# Independently evolved pollution resistance in four killifish populations is largely explained by few variants of large effect

**DOI:** 10.1101/2023.04.07.536079

**Authors:** Jeffrey T. Miller, Bryan W. Clark, Noah M. Reid, Sibel I. Karchner, Jennifer L. Roach, Mark E. Hahn, Diane Nacci, Andrew Whitehead

## Abstract

The genetic architecture of phenotypic traits can affect the mode and tempo of trait evolution. Human-altered environments can impose strong natural selection, where successful evolutionary adaptation requires swift and large phenotypic shifts. In these scenarios, theory predicts the influence of few adaptive variants of large effect, but empirical studies that have revealed the genetic architecture of rapidly evolved phenotypes are rare, especially for populations inhabiting polluted environments. *Fundulus* killifish have repeatedly evolved adaptive resistance to extreme pollution in urban estuaries. Prior studies, including genome scans for signatures of natural selection, have revealed some of the genes and pathways important for evolved pollution resistance, and provide context for the genotype-phenotype association studies reported here. We created multiple quantitative trait locus (QTL) mapping families using progenitors from four different resistant populations, and genetically mapped variation in sensitivity (developmental perturbations) following embryonic exposure to a model toxicant PCB-126. We found that a few large-effect QTL loci accounted for resistance to PCB-mediated developmental toxicity. QTLs harbored candidate genes that govern the regulation of aryl hydrocarbon receptor (AHR) signaling, where some (but not all) of these QTL loci were shared across all populations, and some (but not all) of these loci showed signatures of recent natural selection in the corresponding wild population. Some strong candidate genes for PCB resistance inferred from genome scans in wild populations were identified as QTL, but some key candidate genes were not. We conclude that rapidly evolved resistance to the developmental defects normally caused by PCB-126 is governed by few genes of large effect. However, other aspects of resistance beyond developmental phenotypes may be governed by additional loci, such that comprehensive resistance to PCB-126, and to the mixtures of chemicals that distinguish urban estuaries more broadly, may be more genetically complex.

## INTRODUCTION

Human activities cause rapid and dramatic environmental changes that perturb the health and performance of many wild species. Examples of contemporary environmental change include habitat loss, introduction of invasive species, harvest pressure, climate change, and pollution (Vitousek et al. 1997). These perturbations can cause population declines and extinctions. Adaptive evolutionary change is one mechanism whereby populations may respond and escape local extinction (Palumbi 2001; Smith and Bernatchez 2008; Hoffmann and Sgro 2011; Sih et al. 2011; Hendry et al. 2017). However, specific features of species and of the environment may influence the pace and extent of adaptive phenotypic evolution, and it is important to understand how these features interact with natural selection to predict the likelihood of species’ persistence in the Anthropocene. Relevant features of species include population size, generation time, standing genetic variation, phenotypic plasticity, gene flow, and the genetic architecture underlying relevant phenotypic traits (Barrett and Schluter 2008; Bell and Gonzalez 2009; Bell 2013; Bergland et al. 2014; Orr and Unckless 2014; Kreiner et al. 2018). Relevant features of the environment include the pace of change, the degree of change, and the complexity (dimensionality) of change (Tilman and Lehman 2001; Lindsey et al. 2013; Lourenço et al. 2013; Bay et al. 2017; Whitehead et al. 2017).

The genetic architecture of rapidly evolving traits has emerged as a key area of study. Within this context we consider genetic architecture to include the number of loci, their effect sizes, their interactions, and the ways in which they fit into molecular pathways. It is generally understood that variation in most phenotypic traits is controlled by many loci, most with small effects (Barton and Keightley 2002), and that local adaptation is therefore often polygenic, requiring shifts in allele frequencies at many loci (Pritchard and Di Rienzo 2010). However, the architecture of adaptation in response to anthropogenic change may be atypical. This is because the pace of environmental change is so rapid, and new phenotypic optima may be distant from historical averages. Furthermore, selection coefficients can be very large, such that successful adaptation requires large, rapid phenotypic shifts, potentially favoring fewer loci with much larger effects. Indeed, population genetic theory is pivoting to confront these new scientific challenges (Messer et al. 2016), and simulation studies are examining how the genetic architecture of traits affects the pace of adaptation in changing environments (Jain and Stephan 2017; Stetter et al. 2018). Our goal in this work is to understand the genetic architecture that underlies an example of this rapid adaptation, the evolved resistance to the toxic effects of extreme environmental pollution in the Atlantic killifish (*Fundulus heteroclitus*), and to exploit this system as a model to better understand how adaptation to complex human-altered environments may proceed.

*F. heteroclitus* is a small non-migratory fish inhabiting estuarine habitats of eastern coastal North America. Many estuaries in this region have been heavily impacted by urbanization, which in the 20^th^ century included the release of large quantities of a diversity of highly toxic, persistent, bioaccumulative organic pollutants, including chlorinated dibenzo-*p*- dioxins (“dioxins”), polycyclic aromatic hydrocarbons (PAHs) and polychlorinated biphenyls (PCBs). These pollutants are highly toxic to most vertebrates, even at very low concentrations, where the most sensitive responses to exposure include defects in cardiovascular system development. Nevertheless, populations of *F. heteroclitus* and its sister species *F. grandis* (found from the Atlantic coast of Florida into the Gulf of Mexico) have rapidly and repeatedly evolved resistance to them, even in some of the most intensely polluted sites (Nacci et al. 2010).

Prior work in killifish has shed some light on the genetic architecture of the resistance trait. Evidence has come from comparative transcriptomics (Whitehead et al. 2010, 2012; Oleksiak et al. 2011), genome scans (Williams and Oleksiak 2011; Reid et al. 2016; Oziolor et al. 2019), and association studies (Nacci et al. 2016). Transcriptomic responses to PCB-126 exposure (a model toxicant) showed that global desensitization of the aryl hydrocarbon receptor (AHR) signaling pathway distinguished all resistant from all sensitive populations. This is consistent with other findings from killifish and other species, where AHR signaling mediates toxicity of ubiquitous pollutants such as dioxins and some PCBs and PAHs. Many dioxins, PCBs and PAHs interact directly with the AHR, and the subsequently activated molecular signaling cascade is largely responsible for the toxicity observed in vertebrates, especially fishes during early (embryo-larval) development (Clark et al. 2010; King-Heiden et al. 2012; Shankar et al. 2020). In fact, activation of AHR signaling defines a class of highly toxic, persistent and bioaccumulative pollutants known as ‘dioxin-like compounds’ (DLCs). DLCs typically include dioxins and coplanar PCBs. Though PAHs share some mechanisms of toxicity (such as AHR activation), as a class of chemicals they are typically considered distinct from DLCs.

Association and genome scan studies have also further implicated AHR pathway elements. A microsatellite-based QTL study found that resistance in the New Bedford Harbor (MA) population was associated with a small number of genomic regions, most prominently including a region containing a key AHR signaling gene, aryl hydrocarbon receptor interaction protein (AIP) (Nacci et al. 2016). Genome-wide scans for signatures of natural selection, which included multiple resistant and sensitive populations from both *F. heteroclitus* and *F. grandis*, found loci that are key parts of the AHR signaling pathway to be major targets of selection in populations from polluted sites (Reid et al. 2016). Three loci harbored top-ranked signatures of selection in all four resistant *F. heteroclitus* populations studied – a strong signal of convergent evolution: One locus contained a tandem pair of AHR genes (there are two tandem AHR pairs in the *F. heteroclitus* genome); a second locus contained AIP, a key binding partner of AHR; and a third locus contained Cyp1A, a major downstream regulatory target of AHR. A number of other AHR signaling pathway genes were found to be targets of selection in just three or fewer resistant populations. In *F. grandis,* the same tandem AHR pair was found to be under selection at polluted sites, where the selected variant had introgressed from *F. heteroclitus* (Oziolor et al. 2019). Modeling indicated that selection coefficients for the AIP locus in *F. heteroclitus* and the AHR locus in *F. grandis* ranged from 0.3-0.8. This means that selected variants could rise to fixation from very low frequency on a time scale of tens of generations, which is consistent with the onset of pollution in the mid-20^th^ century.

While these studies have been successful in unveiling some critical aspects of the genetics of resistance (the AHR pathway is important, a handful of loci likely have very strong effects on the resistance phenotype), their correlational nature leaves many important questions unanswered. Genome scans can reveal suites of loci that may contribute to fitness (e.g., by revealing signatures of natural selection), but they do not provide information that directly connects genotype to phenotype. This means that genome scans cannot reveal whether the selected loci all contribute to a single adaptive phenotype or many, nor can they determine the effect size of a locus, if they are redundant, or if they interact. It is likely that multiple traits are simultaneously under selection between different environments (Langerhans 2018). For example, in complex environments like urban estuaries (Rivkin et al. 2019), loci associated with adaptation to urban pathogens may tend to accompany loci associated with urban toxicant resistance. As well, under these intense selective conditions, it may be likely that multiple redundant alleles are favored by selection, resulting in soft selective sweeps (Messer and Petrov 2013). To extend our understanding of the genetic architecture of rapidly evolved pollution resistance in this system requires experimental manipulation.

In the study reported here, we used an experimental approach to further reveal the genetic architecture of adaptation to pollution in *F. heteroclitus.* We created one QTL mapping family for each of four DLC-resistant killifish populations by crossing one killifish from each of four resistant populations with a killifish from a common sensitive population. Embryos from mapping families were exposed to a model toxicant (PCB-126) at a dose known to distinguish sensitive from resistant individuals, and each embryo was scored for developmental abnormalities indicative of their sensitivity to toxicity. We then genotyped ∼48 of the most-resistant and ∼48 of the most-sensitive individuals from each family. Our QTL analysis of developmental abnormalities revealed a small set of loci that accounted for a large portion of phenotypic variation in each population. Some QTL were shared among populations, whereas others were not. Most but not all QTL were in regions that also showed strong signatures of selection in wild resistant fish, and many regions that had very strong signatures selection in resistant populations were not found to be associated with resistance in this experiment. These results highlight the power of integrating quantitative and population genetic approaches to understand the genetic basis of adaptation and represent a large advance in our understanding of the genetic architecture underlying rapid parallel adaptation to extreme pollution in urban killifish.

## METHODS

### Breeding Design

We created mapping families from the same source populations as those described in (Reid et al. 2016), which have been characterized previously for sensitivities to PCB-126 (Nacci et al. 2010). DLC-resistant source populations originated from three sites from the northern portion of the species’ range: New Bedford Harbor, MA, USA (NBH: 41.6676N, 70.9159W), Bridgeport Harbor, CT, USA (BRP: 41.1570N, 73.2189W), Newark, NJ, USA (NEW: 40.7006N, 74.1223W), and one site from the southern portion of their range: the Atlantic Wood site of the Elizabeth River, Virginia, USA (ELR: 36.8078N, 76.2945W) (Figure 1), designated as “T1”, “T2”, “T3”, and “T4”, respectively in (Reid et al. 2016). The DLC-sensitive source population originated from Block Island, RI, USA (BLI: 41.1818N, 71.5793W) and was designated “S1” in (Reid et al. 2016). Husbandry and phenotyping procedures used in this QTL study were similar to, and conducted in the same laboratory as, those used in a previous QTL study (Nacci et al. 2016); in fact, the NBH mapping families used in both studies are derived from the same genetic line (identified as ‘Cross A’ in (Nacci et al. 2016)).

**Figure 1.**
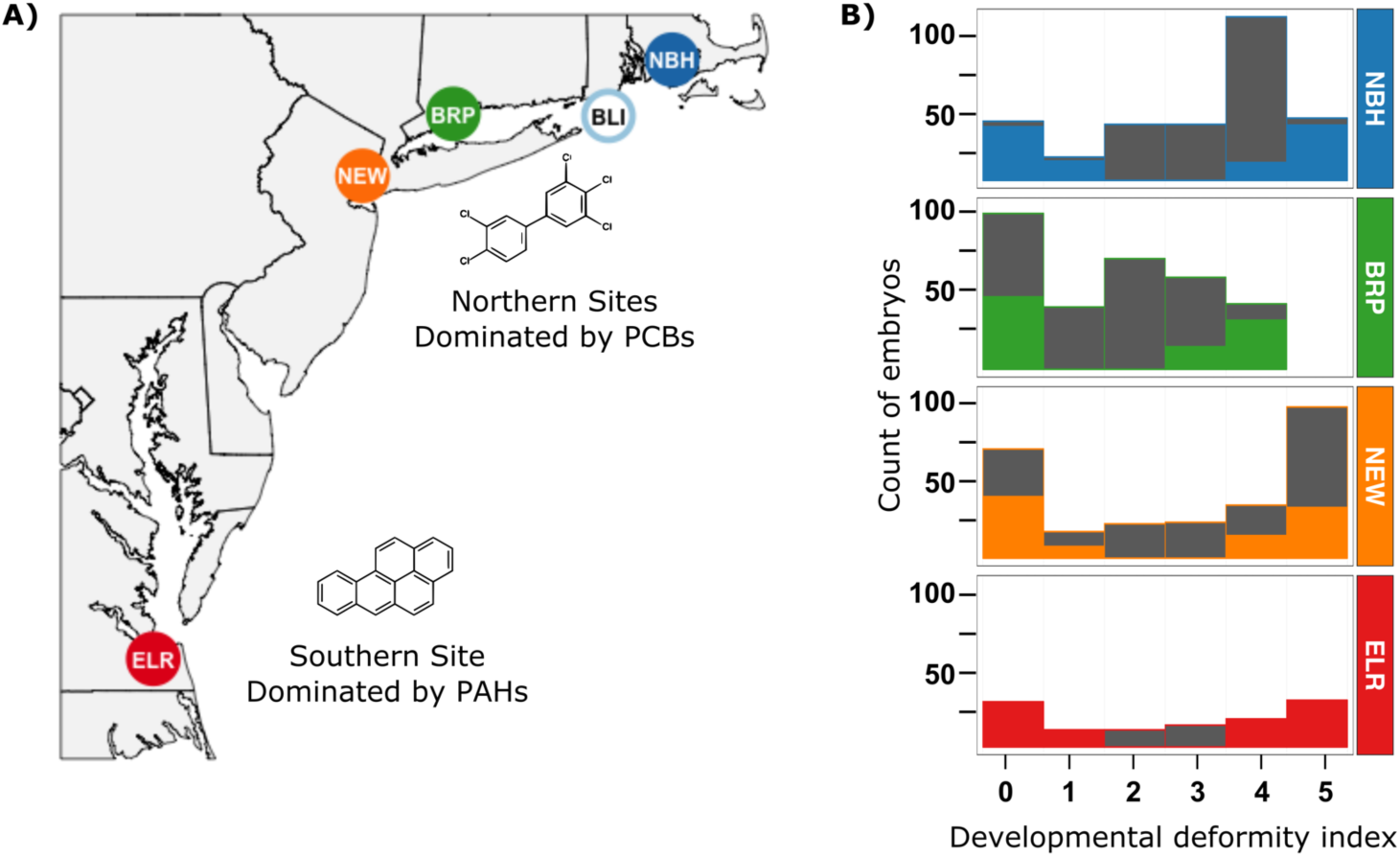
A) Locations of source populations for founders of mapping families. Sites are characterized by complex mixtures of pollutants but dominated by either polychlorinated biphenyls (PCBs) or dioxins in the three Northern sites (New Bedford Harbor, MA (NBH), Bridgeport Harbor, CT (BRP), and Newark, NJ (NEW)), or polycyclic aromatic hydrocarbons (PAHs) in the Southern site (Elizabeth River, VA (ELR)). All polluted site populations (filled circles) have been previously shown to be resistant to PCB-126. Open blue circle is the clean reference site (Block Island, RI (BLI)) and source for one of the progenitors for each of the mapping families. B) Bar graphs show the phenotypic distribution (developmental deformity index, x-axis) of embryos from each of the F2 intercross mapping families at 10 days after an exposure to a discriminating dose, (200 ng PCB-126/L in sea water). Full bars show the total count of embryos collected and assigned a deformity score, which ranged from 0 (no deformities) to 5 (most severe deformities) (the BRP families were scored on a scale where a score of 4 was maximum). Colored portion of each bar indicates the subset of embryos that were selected for genotyping.

We created QTL mapping families using wild killifish that were maintained in the lab (Narragansett EPA), where individuals from BLI (sensitive population) were then bred to individuals from each resistant population (NBH, BRP, NEW, or ELR) to produce F1 hybrid families. F1 hybrid individuals were subsequently paired to produce the F2 mapping families. The specific identities of mapping families were NBH 1105, BRP 812, NEW 1413, and ELR 1433. Over the course of this study, we tried slightly differing breeding strategies to maximize the number if progeny with divergent phenotypes (see Figure S1 for the distributions of phenotypes per family). Thus, breeding designs for mapping families varied slightly. Briefly, the NBH and ELR F2 hybrid mapping families descended from mating one wild-caught female killifish from the resistant population (NBH or ELR) and one wild-caught male killifish from BLI. The NEW mapping family descended from mating one lab-bred female killifish from BLI and one lab-bred male killifish from NEW, while the BRP mapping family descended from two lines, each produced by mating one wild-caught BLI female and one wild-caught BRP male. Thus, resistance was inherited through females in the NBH and ELR families, and through males in the NEW and BRP families. Previous studies have demonstrated that inheritance of resistance is not sex linked (Nacci et al. 1999). Furthermore, there was a single tolerant progenitor for NBH, ELR and NEW families (sib:sib matings), while the BRP family was derived from two tolerant progenitors (non-sib mating).To characterize tolerance, we challenged embryos from each of the four F2 mapping families with a discriminating dose of the model DLC (PCB-126; 3,3’,4,4’,5-pentachlorobiphenyl), scored each for developmental abnormalities associated with DLC toxicity, and then archived them for later genotyping (described below).

### Phenotyping

We exposed and phenotyped embryos as described in (Nacci et al. 2016). Briefly, we exposed embryos from all families to acetone-solubilized PCB-126 dissolved in sea water at a concentration (200 ng/L) that discriminates sensitive from resistant killifish. The exposure lasted from 1 day post-fertilization (dpf) to 7 dpf, followed by transfer to clean seawater. At 10 dpf, we observed the embryos microscopically and scored them for the presence or absence of developmental deformities, where the appearance of multiple abnormalities is tightly correlated with reduced probability of larval survival, and is therefore a good proxy for DLC sensitivity.

Abnormalities typical of DLC-mediated toxicity included pericardial edema, yolk sac edema, heart malformation, tail hemorrhages, body hemorrhages, and abnormal size. Each embryo was scored as the sum of these abnormalities, from 0 (no deformities) to 5 (most deformities), which represents the full range of sensitivity displayed by originating killifish populations. Though the number of abnormalities described above includes 6 phenotypes, the abnormality score was capped at 5 (4 for BRP); in our experience, any individual scored 4 or higher does not survive past two weeks post-hatch. After phenotyping, we flash froze embryos and stored them at -80°C until DNA extraction.

### Genotyping

We used balanced selective genotyping (or two-tail genotyping) (Lebowitz et al. 1987; Lander and Botstein 1989), where we selected ∼48 individuals from each of the most sensitive and most resistant ends of the phenotypic distribution for each family. We used RAD-seq (Miller et al. 2007) for genotyping these individuals from F2 intercross families and mapping family founders. DNA was extracted from frozen embryos with a proteinase-K digestion and the Qiagen DNAeasy kit. Sequencing libraries were prepared for RAD-seq by ligating individual barcodes and NEBnext Illumina oligonucleotides to genomic DNA at *SbfI* cut sites. Four lanes of paired-end (PE-100) sequence data (Illumina HiSeq 2500) were collected (one lane per plate of 96 samples) to enable genotyping of offspring and founders at RAD sites. After de-multiplexing by barcode and evaluating the quality of each sample with FASTQC, sequence reads were aligned to the linkage-mapped *F. heteroclitus* assembly ((Miller et al. 2019); EBI BioStudies accession S-BSST163; this map orders scaffolds from the *F. heteroclitus* reference genome assembly Fundulus_heteroclitus-3.0.2, NCBI BioProject PRJNA177717). We assigned read group information and aligned reads with BWA-MEM using default parameters, marked duplicates with SAMBLASTER, sorted reads with SAMTOOLS, and flagged improperly paired reads with BAMTOOLS (Barnett et al. 2011; Faust and Hall 2014). We used FreeBayes to call genotypes on all populations simultaneously (Garrison 2018). We clustered sample genotypes with metric MDS for QAQC and to cluster genotypes at chromosome 5 to assign sex to the immature embryos (Figure S2) so that sex could be included as a covariate in the QTL search and model fit. Families were each split into separate datasets and filtered for genotypes from repetitive sequence alignments and family-specific invariant sites with PLINK 1.9 (Chang et al. 2015).

Genotypes for each mapping family were formatted to be loaded as independent crosses in R/qtl (Broman et al. 2003). In R/qtl, offspring genotypes were filtered based on the number of genotypes per locus (>12.5% missing data) and the level of segregation distortion (p<0.001) from a 1:2:1 ratio following R/qtl recommendations (Broman and Sen 2009).

Outcrosses in R/qtl are assumed to be from inbred founders, so we limited our analysis to markers that were homozygous for alternate alleles in the founders (double heterozygotes in F1 parents). Our segregation distortion filter was conservative because selective genotyping is expected to lead to some distortion, particularly at QTL of large effect (Xu 2008). After filtration, markers that were genotyped in the founders and F2 embryos were assigned to a consistent allele (A or B) within and among linkage groups. Any markers that we could not confirm with founder genotypes were filtered out so that those remaining were clearly linked with other markers in the linkage group (formLinkageGroups in R/qtl with a recombination frequency < 0.1 and LOD score of 10). We initially anchored markers to their physical position along chromosomes, but then used recombination frequency to reorder (assign a different order than the initial meiotic map) markers and estimated mapping distance in this order with the Kosambi mapping algorithm (Kosambi 1943). Differences from the initial meiotic map may be the result of genome structural variation within a mapping population or read mapping and genome assembly errors. We then checked whether the final filtered marker set sufficiently covered the reference genome physical map and provided a sensible mapping order by visualizing the pairwise recombination frequency and LOD linkage matrix for each chromosome, as well as the relationship between genetic and physical distances along chromosomes (Figure S3). After filtering, the number of linked markers were: 37,304 for the NBH family, 16,725 for the BRP family, 27,119 for the NEW family, and 57,489 for the ELR family.

In addition to genome-wide genotyping with RAD-seq, we also sought to confirm genotypes at two discrete loci: at the loci that encode aryl hydrocarbon receptor 2a/1a (AHR2a/1a) on chromosome 1 and aryl hydrocarbon receptor interacting protein (AIP) on chromosome 2. These loci encode proteins that are core components of the signaling pathway that is adaptively de-sensitized in resistant fish (Whitehead et al. 2012), were implicated as a QTL in the NBH population (Nacci et al. 2016), and show among the strongest signatures of natural selection in resistant populations (Reid et al. 2016). Since the adaptive haplotypes are not fixed in resistant populations (Reid et al. 2016), we sought to confirm whether adaptive haplotypes were segregating in our mapping families. We genotyped ELR offspring with sensitive (scores 4 and 5) and resistant (scores 0 and 1) phenotypes for a deletion that spans the last exon of AHR2a and the first six exons of neighboring AHR1a (99.7 kb deletion; Figure S4) which exists at 81% frequency in wild ELR fish but is absent in sensitive fish from a nearby reference population (methods described in (Reid et al. 2016)). The genotypes at the AHR2a/1a locus (presence/absence of a deletion; Table S1) were added to the QTL analysis with RAD-Tag markers to test for linkage with other chromosomal markers and for an association with the resistant phenotype. Two AIP non-synonymous SNPs at amino acid positions 224 and 252 were genotyped in a subset of mapping family individuals (n=8 for each family, 4 each sensitive/resistant, except for n=35 in ELR for SNP252; Tables S2 and S3) and in the founders of each cross. We amplified a 1.4 kb genomic fragment with PCR primers AIP3F (5’- GGCGCTATACCCGCTCGTGTCC-3’) and AIP5R2 (5’-CTTCATATTTGAAGACGAGGGAGG-3’) using 10 ng genomic DNA and Advantage DNA polymerase (Clontech) with the following cycling conditions: [94°C, 1 min]; [94°C, 5 sec; 68°C, 2 min] 35 X; [68°C, 5 min]. The amplified product was direct-sequenced with the AIP5R2 primer for SNP analysis.

### QTL analysis

Prior to interval mapping, we performed a marker regression on un-filtered markers to set the priors for exploring QTL model-space for interval mapping (Figure S5). We then performed interval mapping on the filtered markers (see above) in R/qtl to test the single QTL model on each chromosome via multiple imputation (n=500) and Haley-Knott regression, which estimates the most likely interval genotypes and QTL position between RAD markers from the log posterior distribution. If the LOD score for the single QTL test exceeded our permuted threshold (top 15% of the null distribution) in a single QTL scan, the position of the highest LOD score was added to the full QTL model. We compared multiple single and full-QTL models between methods (normal, binary, transformed data with parametric and non-parametric models). We examined models that included only the initial two major QTL discovered by marker regression, as well as more inclusive models that included the minor effect QTL that were discovered by interval mapping. QTL were dropped if they did not significantly change the fit of the full-model in the drop-one-qtl analysis. The full model in R/qtl estimated the additive and dominance effect of each of the QTL in the model. We compared their relative effect sizes to illuminate the genetic architecture of resistance in each of the mapping families.

### Selection Scan Data

To further refine our understanding of the genetic variation that contributes to resistance, we tested for overlap between previously identified signatures of selection from population genomics studies (Reid et al. 2016) and the QTL identified here by aligning both datasets to the same chromosomal coordinates. This approach has been effective for connecting QTLs to strong selection from domestication and breeding (Rubin et al. 2012). Signatures of selection were identified from whole-genome scans of diversity and divergence between each of the four resistant populations and nearby non-resistant populations (Reid et al. 2016). In the population genomics studies, genomic scaffolds were scanned for extreme values of pi, Fst, and Tajimas D in 5kb windows to identify signatures of strong recent natural selection. The selection signature thresholds for each statistic were set using a demographic model to simulate a null distribution of each statistic. These thresholds were used to convert the statistics into an aggregate z-score. The Z-scores (which included the width and magnitude of the signature region) formed a composite rank for each signature of selection in each polluted population. Fst, delta pi, and composite ranks on the scaffold level genome assembly were lifted over to a physical position on the killifish chromosomal assembly (Miller et al. 2016, *Fundulus_heteroclitus-3.0.2, NCBI BioProject PRJNA177717)* with liftover using a chain-file generated by ALLMAPS (Hinrichs et al. 2006; Tang et al. 2015). QTL intervals were evaluated against the 50 and 200 top-ranked composite signatures of selection from (Reid et al. 2016).

## RESULTS

### Phenotypes

We observed the full spectrum of sensitive to resistant phenotypes in mapping families, indicating that the founders from each of the populations harbored genetic variation for resistance to PCB-126. Frequency distributions of phenotypic variation for the families that were chosen for mapping (Figure 1B) were representative of phenotype distributions from other F2 intercross families created from these same populations (Figure S1). In our analyses, we grouped individuals with phenotypes of 4 and 5 as sensitive and individuals with phenotypes of 0 and 1 as resistant and modeled the resistance phenotype as binary.

### QTL analysis

We identified a small number of loci that account for a large proportion of phenotypic variation in PCB sensitivity in QTL mapping families from all four resistant populations.

Genome-wide marker regression and QTL interval mapping of RAD-Tag markers identified one or two large-effect QTL in each of the families and these QTL tended to be shared between multiple mapping families (Figure 2A, 2B). A large-effect QTL on CHR18 was shared among all four resistant populations. This CHR18 QTL accounted for a large fraction of resistance variation in each family (11% in NBH, 27% in BRP, 40% in NEW, and 34% in ELR). Another large-effect QTL on CHR2 was shared among the three northern resistant populations. This CHR2 QTL accounted for a large fraction of resistance variation in each northern family (40% in NBH, 22% in BRP, and 18% in NEW). Thus, families from the three northern populations shared the same set of two major-effect QTLs on chromosomes 2 and 18 (CHR2 and CHR18, respectively), despite a different breeding design and phenotypic distribution for the BRP family. Together, this set of two QTLs accounted for a very large fraction of variation in sensitivity to PCB-induced toxicity in northern families (70% in NBH, 53% in BRP, and 55% in NEW). We also found that, at least for the NBH mapping family, the QTL on CHR2 appears to be recessive, and the few heterozygous individuals that are phenotypically resistant carry the resistant genotype on CHR18 (Figure 2C). Parameter estimates for the large-effect QTL models are included in Table S4 and File S1. We detected support for medium to small effect QTL that do not overlap between families. When included in the full model, these additional QTL explain a relatively small proportion of variation (variance explained < 6%, drop-one-qtl analysis estimates). We included parameter estimates for large effect loci with the more inclusive models as shown in Table S4 and File S1.

**Figure 2.**
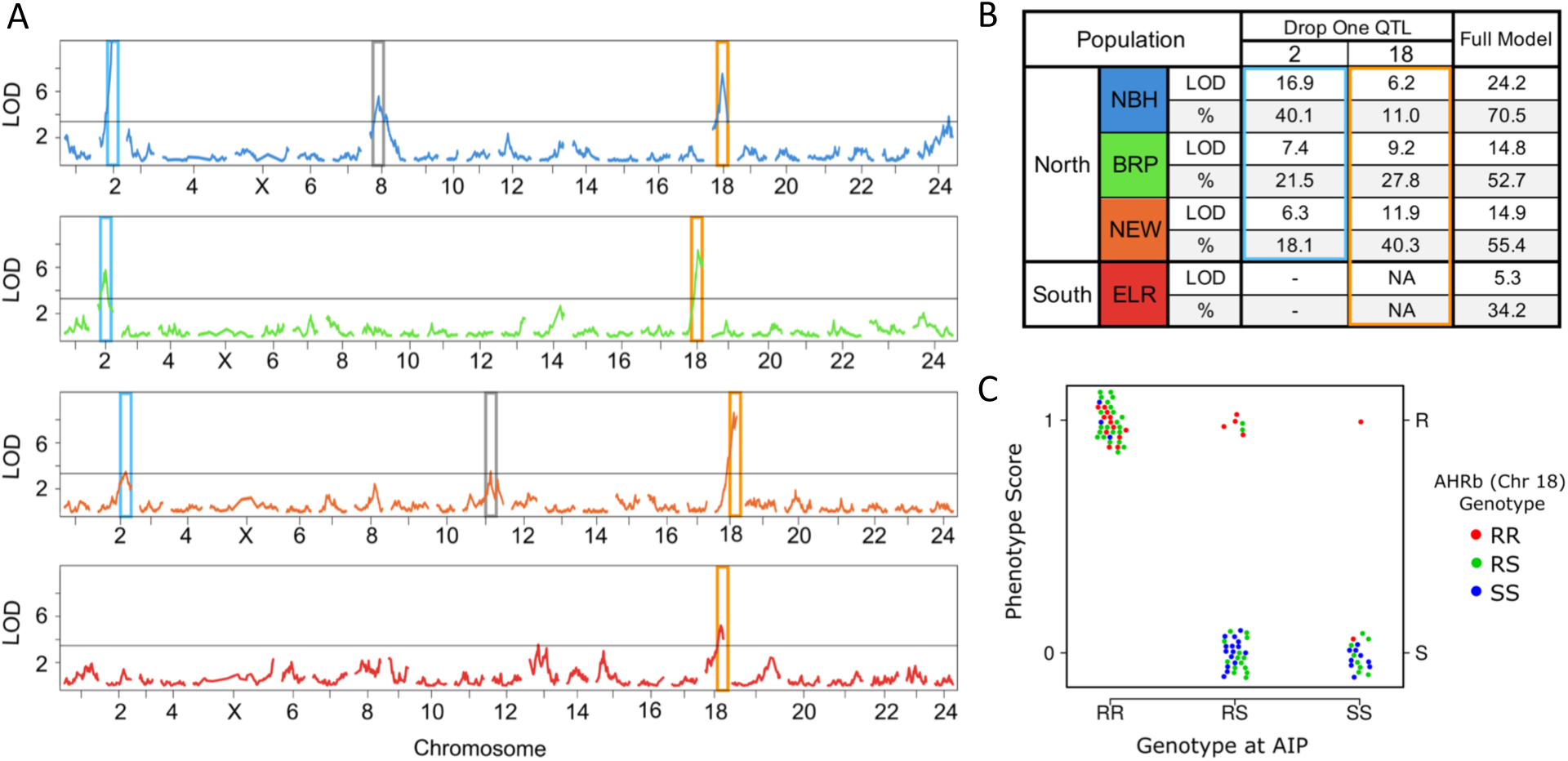
QTL and their contribution to variation in sensitivity to PCB-induced toxicity. A) LOD score results along each chromosome (x-axis numbers) for single QTL scans of RAD-Tag markers in each mapping family. Blue, green, orange, and red series correspond to mapping families from NBH, BRP, NEW, and ELR, respectively. Tall colored boxes (blue and gold) indicate the QTL that fall on the same chromosome/position in multiple families. Grey boxes indicate minor effect QTL that explain a relatively minor proportion of the resistance phenotype and are not shared between populations. Note that the vertical alignment of homologous chromosomes between families is not perfect, because estimated mapping distances for the same chromosome can vary between mapping families because of variation in recombination rates between families and genotyping error. B) Summary of the contribution (drop one QTL analysis) of large-effect QTL (blue and gold boxes) to the full QTL model (LOD and percent variance explained) in each mapping family. C) Genotype-by-phenotype plot for the NBH family at the AIP candidate locus (QTL on chromosome 2) and AHR1b/2b candidate locus (QTL on chromosome 18). The Y-axis groups individuals (points) as either resistant (phenotype malformation scores 0-1 following PCB-126 exposure) or sensitive (phenotype malformation scores 4-5 following PCB-126 exposure). The X-axis distinguishes individuals based on their genotype at the AIP locus (QTL on chromosome 2), where individuals homozygous for the resistant allele, heterozygous, or homozygous for the sensitive allele, are represented by RR, RS, or SS, respectively). The color of points distinguishes individuals based on their genotype at the AHR1b/2b locus (QTL on chromosome 18), where individuals homozygous for the resistant allele, heterozygous, or homozygous for the sensitive allele, are represented by red, green, or blue, respectively). All individuals that carry the RR genotype at the AIP locus are phenotypically resistant (top left group). Those that carry one or no copies of the resistant allele at the AIP locus tend to be phenotypically sensitive (bottom middle and right groups), unless they are also homozygous for the resistant allele at the AHR1b/2b locus (red dots) in which case they tend to be phenotypically resistant (top middle and right groups).

### QTL and Signatures of Selection

We tested for overlap between our QTL peaks and regions of the genome showing signatures of recent strong natural selection from genome-wide scans (Reid et al. 2016), such that overlaps between QTL and selection signatures could narrow the interval for inferring candidate genes. We found that both major QTLs contain genomic regions encoding core components of the AHR signaling pathway. The two large-effect QTL intervals found on CHR2 and CHR18 (Figure 2) include AIP and AHR genes, respectively. As defined by high allele frequency differentiation between resistant and reference populations (Fst) and reduced nucleotide diversity in the resistant population (delta-pi), the AIP genomic region of CHR2 ranks among the strongest signatures of natural selection in all resistant populations (Reid et al. 2016) (Figure 3). This region overlaps with the large-effect QTL for the three northern-population crosses (NBH, BRP, NEW; Figure 3), but not for the ELR mapping family (Figure 2, Figure 3).

**Figure 3.**
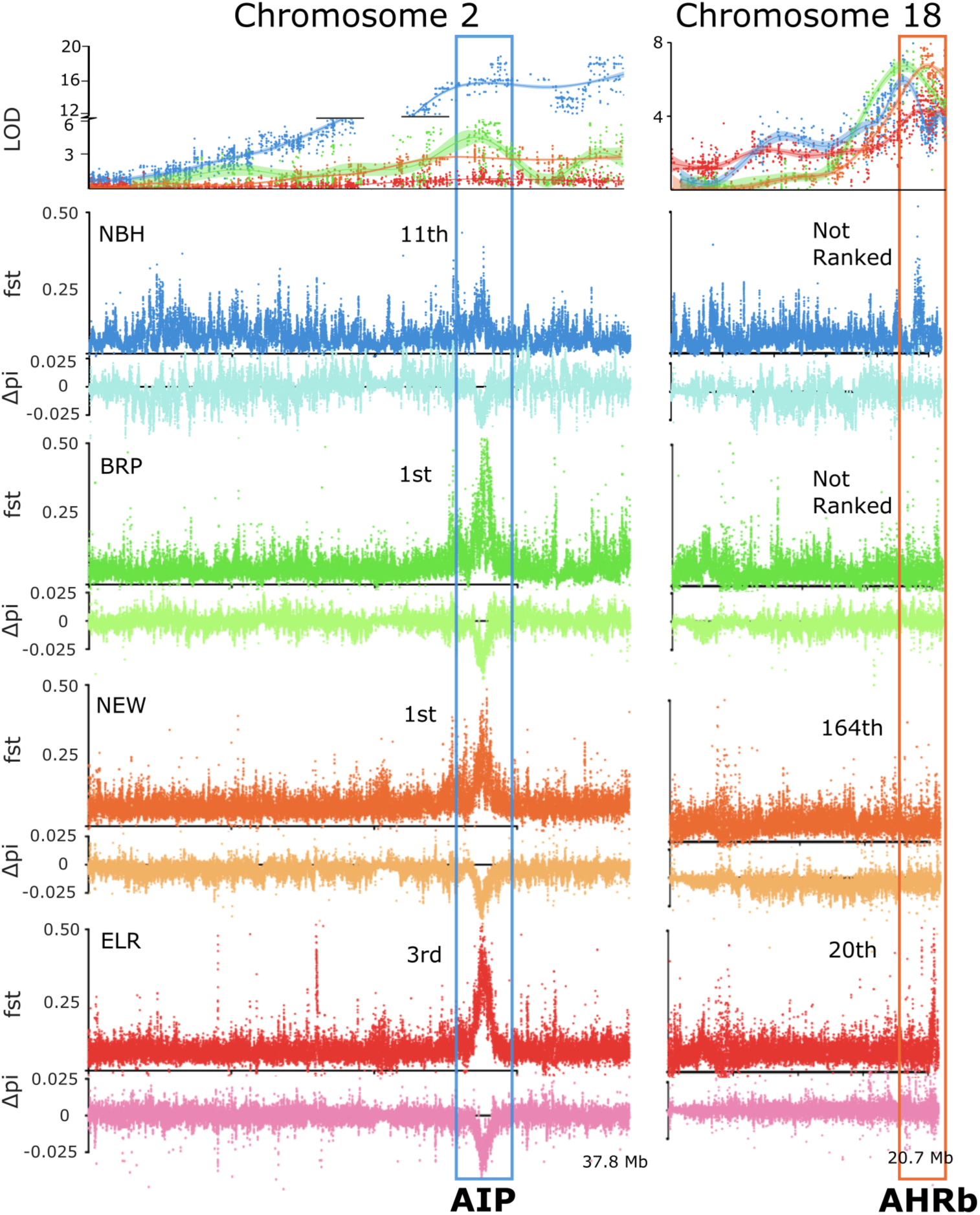
Large-effect QTL (first row) on CHR2 (left column) and CHR18 (right column) aligned with signatures of selection from genome scans for each resistant population (remaining rows). X-axis tick marks represent Mb intervals along the entire length of the two chromosomes. QTL LOD scores (top row) are the results from a single-QTL scan (marker regression) in each of the QTL mapping families (the blue series represents LOD scores from NBH mapping families, green from BRP, orange from NEW, and red from ELR). Panels in the remaining rows include Fst and delta pi between each of the resistant populations and their nearby reference populations (original genome scan data were collected by (Reid et al. 2016)). Elevated Fst and reduced delta pi are signatures of recent strong natural selection. Tall boxes highlight the position and rank of the top-ranked signatures of selection that co-localize to a large-effect QTL. The large effect QTL on CHR2 (which contains AIP) coincides with a highly-ranked selection signature region; though this region shows a strong signature of selection in all four populations, it includes a QTL for only three of the populations (NBH, BRP, and NEW). The large effect QTL on chromosome 18 (which contains AHR1b and AHR2b) is found in all four populations, but coincides with highly-ranked selection signature regions in only two of those populations (ELR and NEW).

The large-effect QTL for all four resistant populations on CHR18 (Figure 2) contains genes AHR1b and AHR2b. This region included the 20^th^ ranked signature of selection in ELR, 164^th^ in NEW, but did not include highly ranked signatures of selection for NBH or BRP (Figure 3).

In addition to the large-effect QTL, we detected support for smaller effect QTL. Markers on chromosomes 8,11,13, and 24 exceeded the permuted threshold in the single QTL scans. However, we found nominal support for the effect and position of these QTL in the full-model with drop-one-qtl analysis. Of the moderate/low effect QTL, the CHR8 QTL had the highest individual LOD score in NBH, but only explained < 2.3% variation when included in the full model. None of these smaller-effect QTL coincide with genomic regions showing strong signatures of selection in the wild resistant populations from which mapping families were derived.

We performed targeted genotyping of the mapping families to verify the presence or absence of putatively adaptive haplotypes that were discovered by the whole-genome scans for selection. In ELR, a highly-ranking signature of recent natural selection on CHR1 was previously discovered to contain a 99.7 kb deletion affecting coding sequences of AHR1a and AHR2a (Figure S4A), and this deletion had swept from low frequency in a nearby reference population (not detected) to high-frequency (81%) in the resistant ELR population (Reid et al. 2016). PCR tests, using primers inside and outside of the deletion region (Figure S4B), verified that the AHR1a/AHR2a deletion on CHR1 is segregating within our ELR mapping family (Table S1).

However, in ELR we detect no evidence for a QTL associated with variable sensitivity to PCB- 126 toxicity anywhere on CHR1 (Figure S5). The PCR genotypes of 40 resistant individuals from our ELR mapping family indicated that eight are homozygous for the deletion, 22 are heterozygous, and ten are homozygous intact. Genotyping of 47 sensitive individuals from our mapping family indicated six homozygous for the deletion, 25 heterozygous, and 16 homozygous intact (Table S1). The phenotype association LOD score of the PCR deletion genotype and RAD markers near the deletion haplotype do not support a QTL on CHR1 for ELR or any of the mapping families (LOD of 0.051 for the deletion genotype in ELR).

Additional targeted genotyping of non-synonymous SNPs in the coding sequence for AIP (CHR2) was used to confirm whether the adaptive genotype at that locus (inferred from results of selection scans from (Reid et al. 2016)) was segregating within our mapping families. We were particularly keen to determine why the ELR mapping family was the only one where the genomic region containing the AIP locus was not a QTL, even though the AIP locus is within one of the highest-ranked signatures of natural selection in all populations including ELR (the AIP locus was within the 3^rd^-highest ranked signature of natural selection detected in ELR; (Reid et al. 2016)). We examined two non-synonymous SNPs in AIP that have high allele frequency differences between resistant populations and their paired sensitive reference populations (Tables S2 and S3). These variants fall within the region containing the adaptive haplotype on CHR2 (Figure 3), and we hereby refer to these variants as “adaptive” insofar as they are indicative of large haplotype frequency differences (and they are in a candidate gene). The TrpèLeu variant in AIP amino acid position 224 (aa224) shows large frequency differences between northern resistant populations (where the Leu residue is between 70-74% frequency) and their paired reference populations (where the Leu residue is between 0-4% frequency) (Table S2). This suggests that the variant is linked to a variant(s) that promotes fitness advantage in polluted estuaries. The Northern resistant founders (NBH, BRP, NEW) of mapping families are all homozygous for the aa224 Leu variant, which is consistent with the support of a QTL at AIP in the Northern mapping families (NBH, BRP, NEW). However, the Trp residue at aa224 is fixed in both the ELR population and its paired reference population (Table S2). In contrast, a different amino acid variant (IleèVal at aa252) shows large allele frequency differentiation at the AIP locus between ELR and its paired reference population, where the Val residue is absent in the reference population, but at 48% frequency in ELR (Table S3). We also found that both the Ile and Val residues were segregating within our ELR mapping family at equal frequency, but variation at this locus was not predictive of resistance (Table S3). This indicates that putatively adaptive variation at the AIP locus was captured in all of our mapping families but was only predictive of resistance to PCB-126 in the three northern populations and not predictive of resistance in the ELR population.

## DISCUSSION

We evaluated the genetic basis for adaptation to a particularly toxic class of pollutants in four populations of killifish that have independently evolved resistance to extreme pollution in urban estuaries of the US Atlantic coast. This study was founded on knowledge that killifish populations from all four polluted sites have converged on a similar level of evolved resistance to the developmental toxicity of a model DLC toxicant, PCB-126 (Nacci et al. 2010).

Furthermore, mechanistic evidence suggested a convergently evolved transcriptional response to PCB-126 exposure in these four populations, which implicated shared adaptive de- sensitization of the AHR signal transduction pathway (Whitehead et al. 2012), which is normally activated by DLC exposure and mediates toxicity (Clark et al. 2010). Subsequent population genetic analyses, including those that specifically examined AHR pathway genes (Proestou et al. 2014; Reitzel et al. 2014) and genome-wide approaches (Reid et al. 2016; Osterberg et al. 2018), revealed AHR pathway genes (among many others) showing signatures of natural selection. An earlier QTL mapping experiment (Nacci et al. 2016) also implicated AHR pathway elements in the NBH population, and provided the foundation for the current studies, where we have now increased the number of genetic markers from hundreds to thousands, and expanded our examination of resistant populations from one to four. We anticipated that genetic variation in AHR pathway elements, and other loci, would be associated with adaptive resistance to PCB- 126 toxicity, but sought to reveal how many loci contributed to resistance, and whether they were shared among families from different populations. We have interpreted results within the context of the comprehensive population genomic study that included all four resistant killifish populations and their nearby sensitive killifish pairs (Reid et al. 2016).

Specifically, we sought to 1) use genotype-phenotype (QTL) mapping to determine the genetic architecture of one component of evolved pollution resistance (developmental toxicity), test whether that architecture is similar or different between our four focal populations, and 3) examine overlap between phenotype-associated loci and loci showing strong signatures of recent natural selection in wild populations. We found that 1) a small number of large-effect QTL loci accounted for resistance to DLC-mediated developmental toxicity, 2) QTLs harbored candidate genes that govern the regulation of AHR signaling, 3) some (but not all) of these QTL loci were shared across all populations, and 4) some (but not all) of these QTL loci showed signatures of recent natural selection in the corresponding wild population (Reid et al. 2016). Reciprocally, some top-ranked genes that were considered strong candidates for PCB resistance inferred from genome scans in wild populations were identified as QTL, but some key candidate genes were not. In what follows, we discuss the nature and implications of each of these key findings.

### Genetic complexity of evolved PCB resistance

Two major large-effect QTL were identified among the four killifish families, with the full model that included these two QTL accounting for a very large proportion of phenotypic variation in developmental sensitivity to our model toxicant (34.2% to 70.1%) (Figure 2). One major QTL was shared among the four mapping families: the QTL on CHR18, which includes candidate genes AHR1b/2b paralogues, accounted for 11% - 43% of the variation in DLC sensitivity. The other major QTL was shared among the Northern resistant killifish but was not identified as a QTL in ELR: the QTL on CHR2, which includes candidate gene AIP, accounted for 18.1 – 45.4% of the variance in DLC sensitivity. A couple of other loci, private to some of the families, accounted for a small fraction of phenotypic variation in some families. For example, minor QTL were noted on CHR8 in the NBH family and on CHR11 in the NEW family. These results are largely congruent with the prior NBH QTL study, where a small number of QTL accounted for 69% of phenotypic variation, with the largest effect loci associated with AHR pathway genes and a region on CHR8 (Nacci et al. 2016). We conclude that variants in a small number of key genes that regulate AHR signaling largely achieve the desensitization of AHR signaling that is protective of DLC-induced development toxicity in urban killifish populations. These results indicate that the genetic basis of evolved variation in developmental sensitivity to PCB-126 is not complex or highly polygenic. Resistance evolved rapidly through increased frequency of large-effect loci, where the adaptive alleles are found at very low frequency (or are undetectable) in sensitive populations from clean habitats (Reid et al. 2016). This core finding is consistent with expectations when adaptation requires a rapid and large shift in the phenotypic optimum (Stetter et al. 2018).

It is possible that variants in AHR and/or AIP may result in similar phenotypic outcomes, by effecting adaptive desensitization of the PCB-induced AHR pathway activation that normally causes developmental toxicity. Variation at the DLC-binding site of the AHR is highly predictive of DLC sensitivity in bird species (e.g., (Farmahin et al. 2013)). Similarly, evolved adaptive DLC resistance in Hudson River (NY, USA) tomcod is largely explained by variants in AHR which affect DLC binding, and that are localized near the site where AIP complexes with AHR (Wirgin et al. 2011). Moreover, AIP is an evolutionarily conserved AHR chaperone that influences the stability of the AHR complex and eventual nuclear translocation of AHR (Bell and Poland 2000; Meyer et al. 2000; Lecoq et al. 2016). AIP loss-of-function mutations in mice protect against some forms of DLC toxicity (Nukaya et al. 2010a). However, the role of AHR-AIP interactions in fish, and whether there are ligand-specific differences in AHR-AIP interactions, are not well understood. In any case, we hypothesize that changes to either locus may result in an altered AHR-AIP-agonist complex that modifies the molecular events that would normally initiate AHR signal transduction.

Few studies have revealed the genomic basis of evolved pollution resistance (but see, e.g., (Wright et al. 2013, 2015)). However, many studies have examined adaptations to environmental pollutants that are toxic by design, such as pesticides, which may offer relevant lessons. Often few loci of major effect are implicated in evolved resistance to pesticides, for example loci that are the biochemical targets of toxicity (e.g., sodium channels that are the molecular targets of neurotoxins), or loci that more broadly govern metabolic detoxification (e.g., cytochrome P450s) (Ffrench-Constant 2013). Though the literature on the genetic basis of pesticide resistance is relatively large, most studies focus on candidate genes; genome-wide examinations of evolved resistance are rare (e.g., (Kreiner et al. 2019, 2021; Van Etten et al. 2020; Pélissié et al. 2022)). Using knowledge from pesticide resistance to make predictions about the genomic basis of pollution resistance may be fraught, partly because genome-wide understanding of the relevant genetics is thin, and partly because the nature of selection in polluted environments may be different from that in agricultural settings. For example, pesticides and antibiotics are typically designed to target very specific biochemical processes; they are usually encountered in high concentrations as single chemicals or simple mixtures. In contrast, multiple toxic pollutants often co-occur in the environment, presenting a highly dimensional fitness challenge (Whitehead et al. 2017). Toxic chemicals are often encountered in human-modified environments where multiple co-occurring habitat alterations may contribute additional selective pressures (e.g., altered competitive interactions, thermal and flow regimes, hypoxia), which further increases the dimensionality of fitness challenges and the required adaptations (Rivkin et al. 2019).

In light of our discovery of few genes of large effect, which is consistent with expectations from the pesticide resistance literature, it is important to consider whether more broadly defined resistance to PCB toxicity, and fitness in polluted urban estuaries more generally, has a simple or complex genetic basis. That is, in this study we only examined resistance to a limited suite of toxic effects caused by a single PCB congener (PCB-126) during a critical developmental period. PCB-126 is only one representative of the many DLCs (albeit a very potent one) and other pollutant classes that may affect fitness throughout the lives of urban fish. We currently do not know whether other aspects of DLC toxicity in embryos or later life stages (e.g., immune dysfunction, endocrine dysfunction, neurological dysfunction, or cancer (White and Birnbaum 2009)) are resolved in resistant fish by the same or different QTL as those identified here. Furthermore, it is plausible that initial costs of adaptive AHR desensitization may have prompted compensatory adaptive fine-tuning to restore some function to AHR signaling and connected pathways. Indeed, sweeps of rare large-effect variants, such as those that we detect, are likely to incur fitness costs, thereby prompting the evolution of cost-alleviating compensatory mutations (Craig MacLean et al. 2010). Consistent with this, our population genomics data indicate signals of selection in other components of the AHR and connected pathways (e.g., hypoxia, cytokine, and estrogen signaling) (Reid et al. 2016; Whitehead et al. 2017). Similarly, our model toxicant PCB-126 is only one of a complex mixture of chemicals that pollute these urban estuaries and threaten fitness (and toxicants are one of a suite of perturbations in urban estuaries (Whitehead et al. 2017)). Populations inhabiting these sites have likely evolved resistance to many of those chemicals (e.g., (Greytak et al. 2010; Bugel et al. 2014; Grans et al. 2015; Celander et al. 2021)). Resistance to some of those chemicals may be underpinned by the same loci as we identified here, but other loci are also likely to contribute to more broadly defined pollution resistance, especially for chemicals with unique mechanisms of toxicity. Indeed, the ELR population has evolved resistance in a site dominated by PAHs, whereas the three northern populations have evolved resistance in PCB-polluted sites. Yet, all four populations are resistant to PCB-126, and we identified one shared major-effect QTL (Figures 2, 3). At least part of the explanation for this is that some PCBs and PAHs share a common mechanism of developmental toxicity mediated through AHR signaling (Billiard et al. 2008; Clark et al. 2010; Incardona 2017). By extension of the above reasoning, the many environmental changes that distinguish urban from natural estuaries include diverse non-chemical perturbations (e.g., biotic interactions, flow regimes, pathogens, etc.) that may drive adaptation through entirely unique suites of traits, pathways, and genes. We conclude that evolved resistance to the developmental toxicity induced by PCB-126 has a relatively simple genetic basis in resistant individuals, while acknowledging that the genetic changes required for adaptation to urban estuaries are likely much more complex.

### Variation in QTL among resistant populations

Our mapping families derived from four focal populations share some similarities and also some differences in the loci that contribute to resistance. Mapping families from the three northern populations share each of the two major-effect QTL (on CHR2 and CH18), whereas the mapping family from the southern population (ELR) is most different and shares just one QTL (on CHR18) with those from the northern populations. Similarities and differences between QTL for mapping families representing different populations may be driven by any of three factors: 1) presence of functionally redundant variants, 2) differences in genetic background, or 2) differences between selective environments; we consider these in sequence.

First, for many adaptations, the genetic basis is polygenic. This could manifest as many variants, each of small effect, contributing to phenotypic differences between two individuals (one resistant, one sensitive), which appears to not be the case for our data presented here.

Alternatively, polygenic adaptation could manifest as multiple redundant large-effect variants contributing to adaptive fitness among different pairs of individuals. That is, multiple different but functionally redundant variants may be segregating within a population that each increase in frequency under selection (e.g., (Goldstein and Holsinger 1992)), such that QTL for mapping families derived from two different resistant individuals could implicate different loci. Indeed, there was limited overlap in PCB resistance QTL among the three mapping families created with different NBH progenitors (Nacci et al. 2016). This type of polygenic adaptation, perhaps better termed “oligogenic”, is likely relevant for toxicant-resistant killifish populations, insofar as the resistance phenotype is fixed but almost no SNPs are fixed in those populations (Reid et al. 2016). Therefore, individuals selected as QTL mapping family progenitors may harbor different combinations of adaptive loci (Nacci et al. 2016). If adaptation is polygenic or oligogenic, then QTL studies (because they include only one or very few progenitors) may not reveal the complete sets of loci that contribute to evolved phenotypes. However, we do not consider this a good explanation for the QTL differences on CHR2 presented here; variants on CHR2 have been subject to strong recent natural selection in all four populations (Figure 3; (Reid et al. 2016; Osterberg et al. 2018)), yet in the present study variation in this region was only associated with resistance in mapping families from NBH, BRP, and NEW populations, but not from ELR. Perhaps the adaptive haplotype was missing from the ELR individual that we selected as the mapping family progenitor? We consider this unlikely since our additional sequencing verified that both putatively adaptive (variant with high frequency in the resistant population) and sensitive genotypes at this locus were segregating in the ELR mapping family (Table S3). However, it is still possible that these variants are not closely linked with the adaptive variant(s). Ongoing high-density genetic mapping should resolve this.

The second factor that could contribute to inter-family variation in QTL is genetic background. The three northern populations are genetically quite similar to each other (average pairwise Fst = 0.09), whereas the southern population has a very divergent genetic background (average pairwise Fst between ELR and each of the northern populations = 0.19) (Reid et al. 2016). Accordingly, the genetic variation in northern populations that pre-dated pollution, and upon which post-pollution selection acted, was likely shared, but the genetic variation available for selection in the southern population was quite different, such that the genotypes underlying parallel adaptation were shared in the north but different in the south. However, we do not consider this sufficient to explain the QTL differences presented here. Although the adaptive haplotypes were different between northern and southern populations, the genomic region on CHR2 (which coincides with the CHR2 QTL) was among the very top-ranked signatures of natural selection in all four resistant populations (Figure 3; (Reid et al. 2016; Osterberg et al. 2018)). If this adaptive variation supported resistance to PCBs then the region on CHR2 should be a QTL in all four populations.

The third factor that could contribute to inter-family variation in QTLs is differences between the environments that select for variants at different adaptive loci, thereby altering genotype-phenotype associations even for parallel-evolved traits. The northern populations are united by halogenated DLCs (PCBs and TCDD) being the dominant toxicants in those sites, whereas the southern ELR site is dominated by PAHs. Though PCBs, TCDD, and PAHs may exert some aspects of developmental toxicity in part through a common mechanism (AHR signaling; (Clark et al. 2010)), the detailed mechanisms of action of these two chemical classes differ (Billiard et al. 2008; Denison et al. 2011a; Incardona 2017). One important difference is that PAHs are readily metabolized by AHR-regulated proteins (potentially creating many reactive metabolites), whereas PCBs are not. Another difference is that dioxins and dioxin-like PCBs mediate cardiac toxicity largely through the AHR, but many PAHs exert their cardiac toxicity through AHR and other mechanisms (Incardona et al. 2005) such as AHR-independent disruption of excitation-contraction coupling (Brette et al. 2014). Clearly, variation at CHR2 has been subject to strong selection in all four populations (Figure 3). Perhaps PCB-induced selection in northern estuaries favored variation at CHR2 that enables PCB-resistant phenotypes, whereas PAH-induced selection in ELR also favored variation at CHR2 but those variants promote aspects of adaptation (e.g., PAH resistance) that are independent of PCB resistance. We consider this a plausible explanation for our data. Ongoing experiments are testing for QTL for multiple classes of chemical resistance (including PAHs) in multiple killifish individuals and populations.

### Coincidence between QTL and signatures of natural selection

Some, but not all, QTLs for PCB resistance showed signatures of selection in wild populations. The QTL on CHR2 in mapping families from NBH, BRP, and NEW overlaps with a top-ranked signature of selection in genome scans of those same populations. This locus harbors the AIP gene, which encodes a key regulator of AHR signaling that is adaptively desensitized in resistant killifish populations. In contrast, the QTL on CHR18 found in all four mapping families was a ranked signature of selection in only two of the four wild populations; this locus was ranked 20^th^ and 164^th^ in ELR and NEW populations, respectively, but no evidence for recent strong selection was detected in genome scans of BRP and NBH populations (Figure 3). One possible explanation is that multiple functionally redundant adaptive variants at this locus contributed to resistance in wild killifish populations, or a single adaptive variant embedded in multiple genetic backgrounds contributed to resistance, and increased in frequency during adaptation. The soft sweeps or sets of partial sweeps that emerge to underlie adaptation (Pritchard et al. 2010) are notoriously hard to detect because of diminished signal of linked selection, though they may underlie much of the adaptive divergence observed among natural populations. Additional studies that, for example, take advantage of linkage-disequilibrium based tests (Pennings and Hermisson 2006) would be necessary to detect a soft sweep on CHR18 and test this hypothesis.

Some, but not all, of the top-ranked signatures of selection in wild populations were associated with PCB resistance. Genome scans that compare wild populations identify loci that affect fitness between environments, but do not directly implicate the agent of selection. Urban estuaries likely pose many fitness challenges for killifish, including chemical pollution, but also changes in biotic interactions and in nutrient, temperature, hypoxia, and flow regimes. The full suite of adaptive solutions to these complex and multifarious fitness challenges can imprint in genome scans. It is therefore not surprising that many loci showing signatures of recent natural selection are not included among the QTL that are associated with resistance to developmental toxicity of one toxicant. However, we were surprised that a large deletion that spans AHR2a and AHR1a on CHR1 (Figure S4A) was not associated with PCB resistance in the ELR mapping family. This is surprising because AHR desensitization is a hallmark of evolved resistance in killifish (Nacci et al. 2010; Whitehead et al. 2010, 2012; Oleksiak et al. 2011; Reid et al. 2016), AHR knockout/knockdown is protective of PCB (or dioxin) toxicity in rodents (Fernandez-Salguero et al. 1996), zebrafish (Prasch et al. 2003; Goodale et al. 2012; Garcia et al. 2018; Souder and Gorelick 2019), and killifish (Clark et al. 2010), and an AHR2a deletion had swept to high frequency in resistant populations of Atlantic tomcod (Wirgin et al. 2011), Gulf killifish (Oziolor et al. 2019), and the ELR population of Atlantic killifish (Reid et al. 2016), and was also found at elevated frequency in NBH and NEW (Reid et al. 2016). Since the deletion was segregating in the ELR mapping family (Table S1), we conclude that it must be associated with some other aspect of evolved pollution resistance, perhaps associated with evolved resistance to PAH toxicity that is independent of the mechanisms that drive PCB toxicity.

One locus that showed among the strongest signatures of selection in wild populations was strongly associated with PCB resistance; the locus on CHR2 harbors the gene that encodes AIP and was the top ranked signature of selection in BRP and NEW populations, third ranked signature in ELR, and 11^th^ ranked signature in NBH. AIP interacts with the AHR and influences DLC toxicity in rodent models (Nukaya et al. 2010b; Denison et al. 2011b). Adaptive variants at this locus were segregating in all four mapping families (Tables S2 and S3), and this was a major-effect locus in NBH, BRP, and NEW mapping families, but was not a QTL in the ELR family (Figure 2). We conclude that strong selection at AIP confers resistance to PCB toxicity in northern populations, but selection at this locus contributes to other aspects of adaptation in the ELR population. Since PAH pollution distinguishes the ELR site from the others, and AHR signaling is important for PAH toxicity, perhaps variation at AIP is important for evolved resistance to PAH toxicity that is independent of the mechanisms that drive resistance to PCB toxicity.

Conspicuously, the QTL region on CHR18 (and the coincident signature of selection region in ELR and NEW) harbors the two killifish AHR paralogs AHR1b and AHR2b (teleost fish typically have 4 AHR paralogs as a result of tandem and whole-genome duplications (e.g. Figure S4A), though some species such as zebrafish have lost one or more copies (Karchner et al. 2005)). The two other AHR paralogs AHR1a and AHR2a are found on CHR1; this locus is under strong selection in urban killifish populations (Proestou et al. 2014; Reitzel et al. 2014; Reid et al. 2016) and was ranked as the 12^th^, 2^nd^, 2^nd^, and 5^th^ strongest signatures of selection in NBH, BRP, NEW, and ELR populations, respectively (Reid et al. 2016). However, variation at AHR1a/2a on CHR1 was not associated with PCB resistance in any of our mapping families.

This was unexpected because AHR2a was shown to mediate developmental toxicity of PCBs and PAHs in killifish (Clark et al. 2010). However, the zebrafish ortholog of killifish AHR2b mediates the developmental toxicity of PCB-like chemicals in that species (Prasch et al. 2003; Goodale et al. 2012; Garcia et al. 2018; Souder and Gorelick 2019), which along with our data suggests a role for the AHR1b/2b locus in fish sensitivity to PCB-induced developmental toxicity. Zebrafish AHR1b protein also exhibits high-affinity binding of DLCs (Karchner et al. 2005), but loss-of-function studies have shown that AHR1a and AHR1b do not mediate the most prominent forms of DLC developmental toxicity in zebrafish (Garcia et al. 2018; Souder and Gorelick 2019). The relative roles of each of the four killifish AHR paralogs are not yet well understood. All four AHR proteins exhibit high-affinity binding to [^3^H]TCDD *in vitro* and TCDD-inducible transcriptional activation in cell culture (Karchner, Franks, Hahn, unpublished results), suggesting that DLC effects in killifish could involve multiple AHRs. Evidence for strong selection at AHR1a/2a but no QTL for PCB resistance at that locus, and QTL for PCB resistance at AHR1b/2b with evidence for weaker and perhaps polygenic selection at that locus, suggests that evolved pollution resistance is underlain by complex interactions among AHR paralogs and their protein partners (e.g., AIP). In any case, it is evident from both genotype-phenotype mapping and scans for signatures of recent natural selection that genetic variation in core regulatory components of the AHR signaling pathways contributes to evolved resistance to pollution in urban estuaries.

### Caveats and considerations

Selective genotyping may have limited power to detect epistatic or small effect loci. We chose selective genotyping because of the power and efficiency (costs savings) that are afforded (Lander and Botstein 1989). However, modeling studies suggest that if there are loci of large effect then the effectiveness of selective genotyping becomes unpredictable (Sen et al. 2009). Clearly, our results reveal loci of large effect, but small effect loci and non-additive or epistatic loci may also be present (e,g., as in (Nacci et al. 2016)). Indeed, our data suggest that the QTL on CHR2 is recessive (Figure 2C). Ongoing studies by our group involve genotyping and phenotyping entire families that span the entire range of sensitive to resistant phenotypes, which should resolve this potential issue.

Although our results indicate that resistance to PCB-126-induced developmental toxicity is underlain by few variants of large effect, one should be cautious in concluding that this evolved resistance is not more polygenic. This is because our experiments included only one resistant individual as the progenitor for each mapping family (except for BRP which had two resistant progenitors), so sampling of within-population variation in resistance was far from thorough. By sampling many more individuals, GWAS studies could reveal more loci that contribute to resistance, especially if resistance is underlain by multiple redundant variants that are not fixed within resistant populations. For example, GWAS for evolved herbicide resistance has revealed many loci, even when large-effect loci were present, and these additional loci were segregating across a range of allele frequencies, suggesting a complex and polygenic architecture (Kreiner et al. 2021). For killifish, some QTL varied between three NBH mapping families in a previous study (Nacci et al. 2016). However, our three mapping families derived from the three resistant northern populations (NBH, BRP, and NEW) all implicated the same major-effect QTL (Figure 2). It is plausible that minor-effect and epistatic QTL, which we had little power to detect in our study, may be variable among individuals, so that the complexity of resistance to PCB-126-induced developmental toxicity during development may be richer than it currently appears.

One should also be cautious about concluding that all aspects of resistance to PCB-126, or adaptation to pollution more broadly, or adaptation to urban estuaries even more broadly, is not highly polygenic. This is simply because multiple traits, each with a simple or complex genetic basis, likely contribute to fitness in these highly altered urban environments (as we discuss in more detail in the section above “*Genetic complexity of evolved PCB resistance”*).

Additional quantitative genetic studies are needed to draw genotype-phenotype associations for the diversity of phenotypes induced by exposure to the array of stressors that distinguish urban estuaries.

## DATA AVAILABILITY

A GitHub repository including scripts for the analyses presented here can be found at https://github.com/WhiteheadLab/killifish-RADseq-4popQTL. Sequence reads have been deposited at NCBI, and are linked with the NCBI BioProject Accession PRJNA944691. Raw phenotypic abnormality scores for individual embryos are included in Appendix S1.

## ACKNOWLEDGMENTS

This research was supported in part by funding from the U.S. National Science Foundation (OCE-1314567 and DEB-1265282 to AW) and the U.S. National Institutes of Environmental Health Sciences (R01ES032323 to AW and MEH, R01ES021934 to AW; R01ES033888 to MEH) and the Superfund Research Program (P42ES007381 to MEH). This publication was made possible by an NIEHS-funded predoctoral fellowship to JTM (T32 ES007059). Its contents are solely the responsibility of the authors and do not necessarily represent the official views of the NIEHS or NIH. Approval does not signify that the contents necessarily reflect the views and policies of the US EPA. Mention of names, products, or services should not be interpreted as conveying official US EPA approval, endorsement, or recommendation.

## Supplemental Figures and Tables

**Figure S1:**
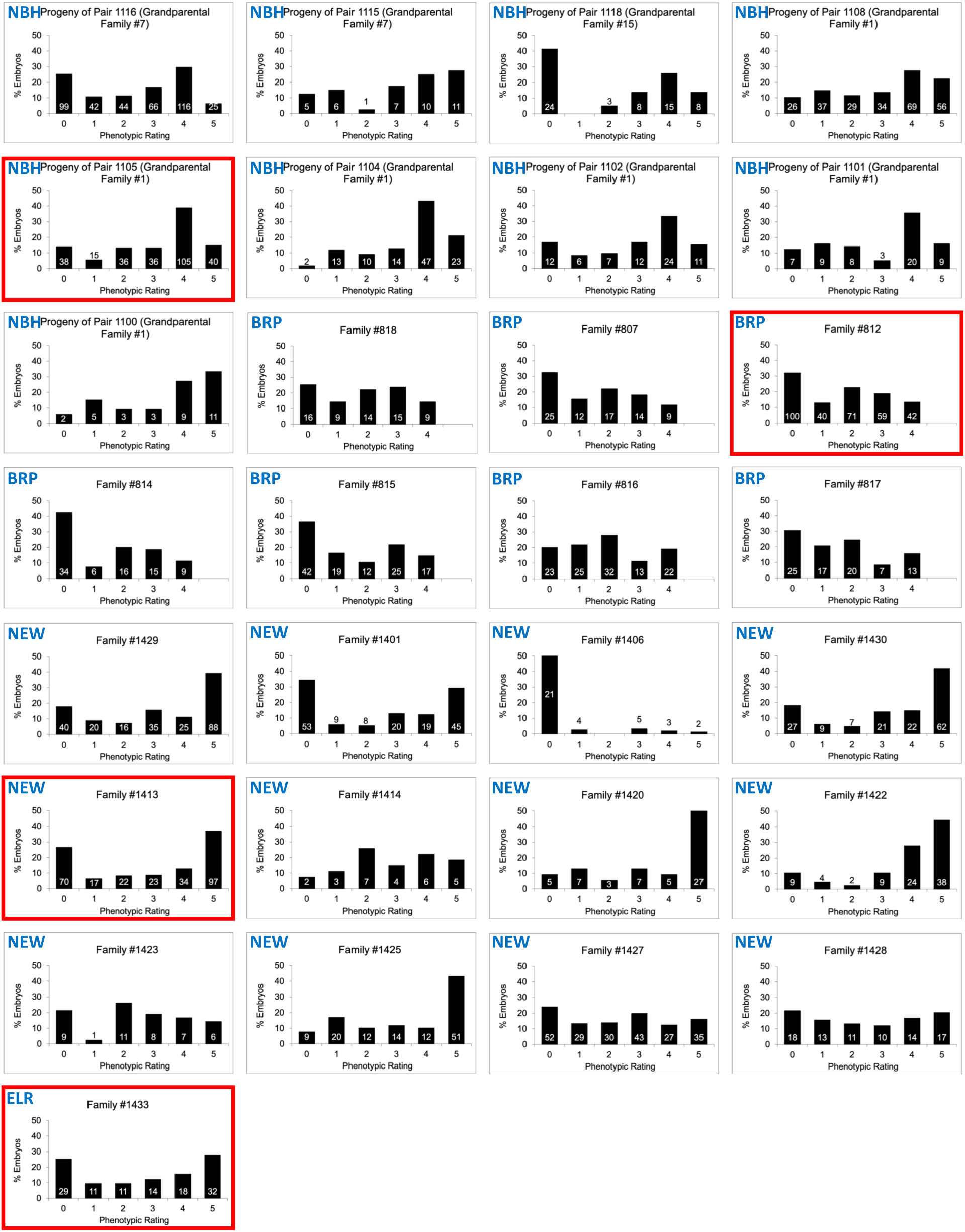
Distributions of embryonic phenotypic sensitivity scores following exposures to PCB-126 for many F2 intercross families created from each of the resistant (NBH, BRP, NEW, and ELR) populations. Individuals were scored on a scale ranging from 0 to 5 for all families except for those from BRP which were scored on a scale ranging from 0 to 4. Families selected for genotyping in the experiments reported here are highlighted in red. Proportion of embryos within each score category is indicated by black bars, where numbers associated with bars represent the number of embryos from that family with that score. Very small families (e.g., <20 individuals) were excluded.

**Figure S2.**
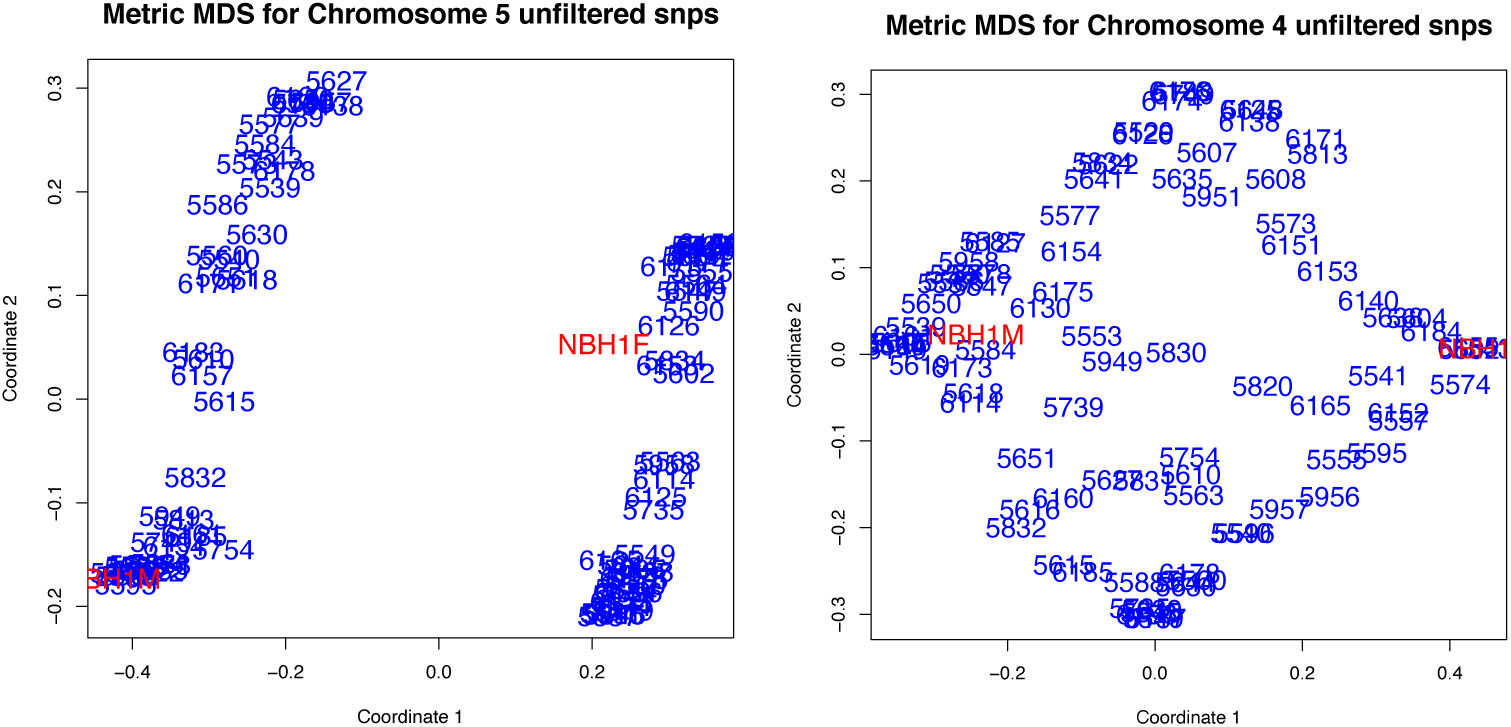
Chromosome 5 is a sex chromosome. Metric MDS plots of unfiltered markers from offspring (blue) and founders (red, NBH1F = founding female, NBH1M = founding male). For the putative sex chromosome (chromosome 5, left), individuals are clearly distinguished by coordinate 1, and clustered with either the male founder (left) or female founder (right). Offspring were assigned the sex of the founder with which they clustered on chromosome 5 (males on the left, females on the right). In contrast, for autosomes (e.g., chromosome 4, right panel), individuals are not clearly distinguished along coordinate 1 nor clustered with founders. Note that chromosomes 5 and 4 as designated here and in (Miller et al. 2019) are designated chromosomes 4 and 23, respectively, in the MU-UCD_Fhet_4.1 assembly.

**Figure S3.**
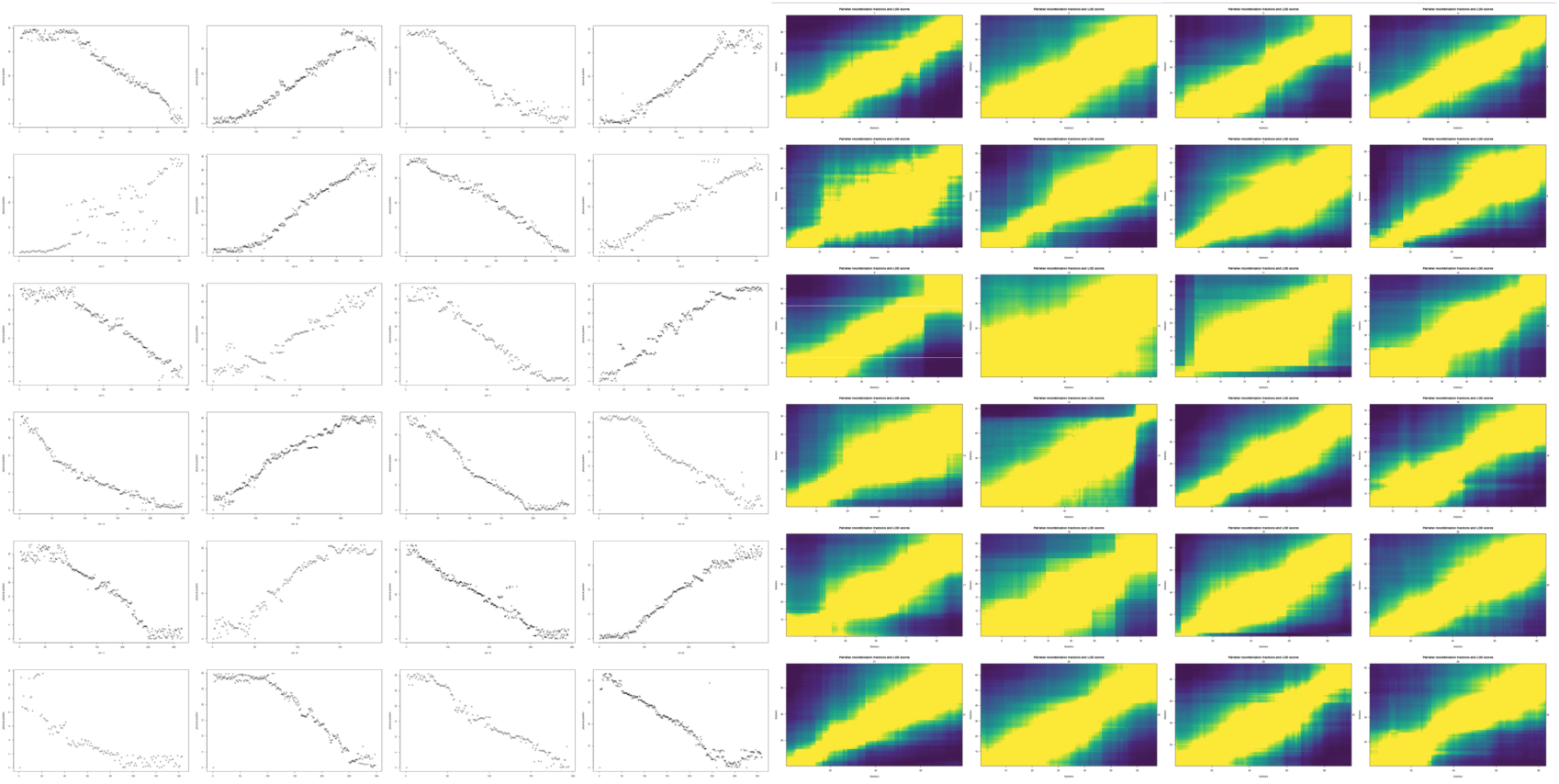
Physical position and linkage of reordered markers for each chromosome (chromosomes 1-24 ordered from top left to bottom right for each panel). Filtered markers were reordered in each mapping family to account for differences from the reference genome genetic map. Left panel: To confirm that the final marker set contained makers along the entire chromosome, we plotted the reference genome physical position (y-axis) of markers in their new order (x-axis). Right panel: Visualizations of linkage (marker pairwise recombination frequency and LOD score) for markers in their new positions confirmed sensible ordering on each chromosome. Note that the chromosome numbering that we refer to in this manuscript is different from the chromosome numbering assigned to the F. heteroclitus genome assembly MU-UCD_Fhet_4.1 (GCF_011125445.2). The chromosome numbering that we use in this manuscript is based on that assigned by Miller et al. (Miller et al. 2019). For key loci discussed in this manuscript, AIP, AHR2a/1a, and AHR2b/1b are localized on chromosomes 2, 1, and 18, respectively, where homologous chromosomes in the MU-UCD_Fhet_4.1 assembly are numbered as 5, 7, and 24, respectively.

**Figure S4.**
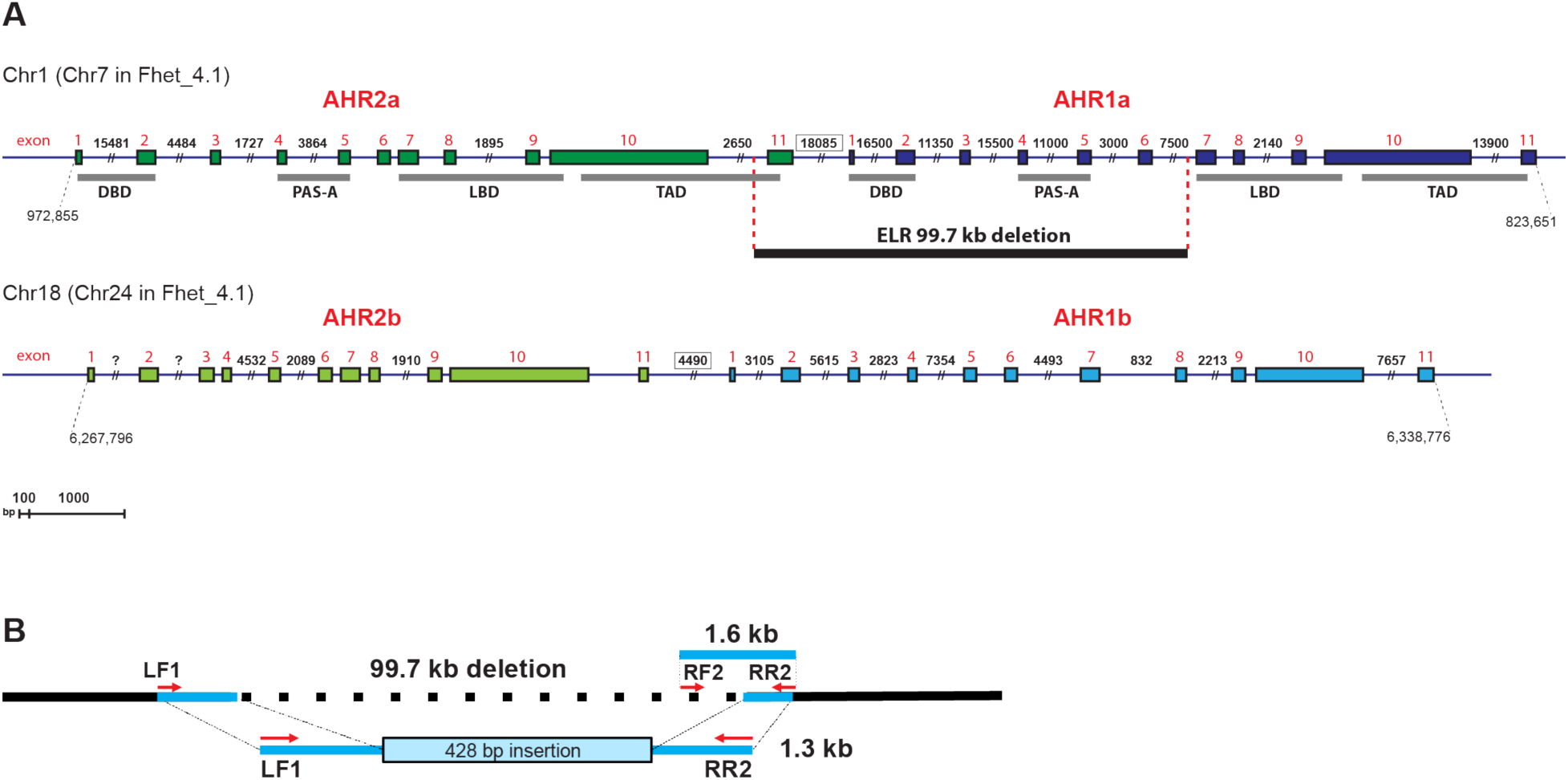
Four paralogs of AHR are found within the *F. heteroclitus* genome. A) Gene models for the four *F. heteroclitus* aryl hydrocarbon receptor (AHR) genes as determined from the Fhet_4.1 genome assembly. Green and blue boxes denote exons. AHR functional domains DBD (DNA-binding domain), PAS-A (Per-Arnt-Sim-A), LBD (ligand-binding domain), and TAD (transcriptional activation domain) are shown with grey bars. The 99.7 kb deleted region of the AHR2a-AHR1a genes in the ELR population is depicted as a black bar below the gene model. Note that chromosome numbering for locations of AHR2a/1a and AHR2b/1b differ between those reported here (based on the numbering assigned by (Miller et al. 2019)), and chromosome numbering for the MU-UCD_Fhet_4.1 assembly (GCF_011125445.2). Chromosome 1 as designated here and in (Miller et al. 2019) is designated chromosome 7 in the MU-UCD_Fhet_4.1 assembly, and chromosome 18 as designated here and in (Miller et al. 2019) is designated chromosome 24 in the MU-UCD_Fhet_4.1 assembly. B) Strategy for confirmation of the ELR deletion spanning AHR2a and AHR1a by PCR. Genomic DNA samples were amplified with primers flanking the left and right junctions of the deleted region (LF1/RR2), as well as within the deletion (RF2/RR2). Primers straddling the deletion (LF1/RR2) resulted in a 1.3 kb PCR fragment in fish heterozygous or homozygous for the deletion, whereas no PCR product was observed in the absence of the deletion. The sequence of the 1.3 kb PCR product matched the genomic sequence flanking the deleted region, except for a 428 bp insertion (Reid et al. 2016). For the primer pair in which one is within the deletion (RF2/RR2), the PCR product was 1.6 kb, whereas no PCR product was observed in fish homozygous for the deletion. (PCR products are not to scale.) The AHR2a-AHR1a deletion in the ELR population was originally characterized as an 83-kb deletion (Reid et al. 2016). The more recent Fhet_4.1 genome assembly reveals that the deletion is actually 99.7 kb.

**Figure S5.**
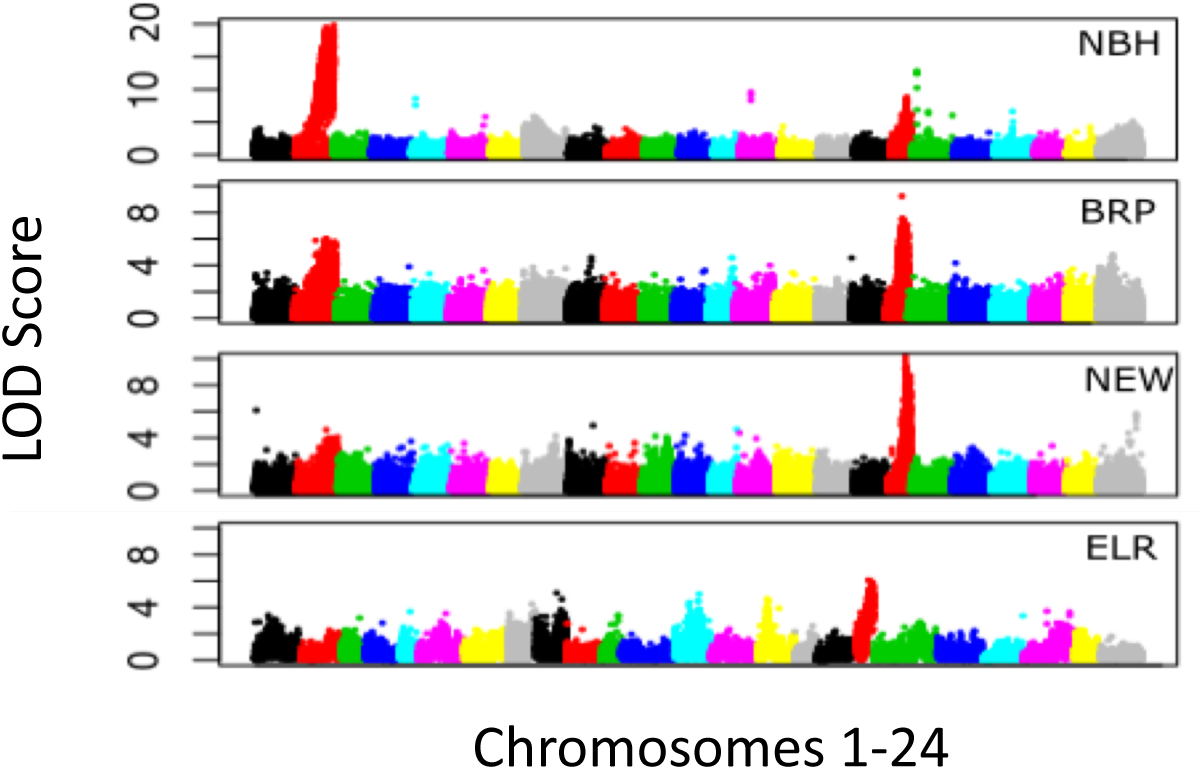
LOD score (y-axis) of genotype by phenotype regression (binary trait) of unfiltered RAD-TAG genotypes for each population. Unfiltered genotypes mapped to physical locations indicate a common locus in all four populations. Different colors indicate different chromosomes, from chromosome 1 on the left to chromosome 24 on the right.

**Table S1.** PCR-verified AHR1a/2a deletion genotype (del = AHRa deletion, wt= no deletion wild type) counts for resistant (phenotype score 0-1) and sensitive (phenotype score 4-5) embryos included in the QTL analysis from the ELR population. The deletion haplotype was segregating in our mapping family, but did not account for any variation in phenotype.

| Phenotype score | Genotype Count |  |  |
| --- | --- | --- | --- |
|  | wt/wt | wt/del | del/del |
| 0-1 | 10 | 22 | 8 |
| 4-5 | 16 | 25 | 6 |

**Table S2.** Mapping family genotype counts and estimated population frequency of the amino acid variant TrpèLeu at residue 224 in AIP.

| AIP (aa 224) |  |  | Count |  |  | Estimated pop. Frequency (Trp/Leu) |  |
| --- | --- | --- | --- | --- | --- | --- | --- |
|  |  |  | Trp/Trp | Trp/Leu | Leu/Leu | Reference | Resistant |
| Phenotype score (0=resistant, 5=sensitive) | NBH | 0 | 0 | 0 | 4 | 1.00/0.00 | 0.26/0.74 |
|  |  | 5 | 2 | 2 | 0 |  |  |
|  | BRP | 0 | 0 | 2 | 2 | 0.97/0.03 | 0.30/0.70 |
|  |  | 5 | 2 | 2 | 0 |  |  |
|  | NEW | 0 | 0 | 2 | 1 | 0.96/0.04 | 0.28/0.72 |
|  |  | 5 | 0 | 1 | 0 |  |  |
|  | ELR | 0 | 4 | 0 | 0 | 1.00/0.00 | 1.00/0.00 |
|  |  | 5 | 3 | 0 | 0 |  |  |

**Table S3.** ELR mapping family genotype counts and estimated population frequency of the amino acid variant IleèVal at residue 252 in AIP.

| AIP (aa 252) |  |  | Count |  |  | Estimated pop. Frequency (Ile/Val) |  |
| --- | --- | --- | --- | --- | --- | --- | --- |
|  |  |  | Ile/Ile | Ile/Val | Val/Val | Reference | Resistant |
| Phenotype | ELR | 0 | 2 | 12 | 5 | 1.00/0.00 | 0.52/0.48 |
|  |  | 5 | 6 | 7 | 3 |  |  |

**Table S4.**
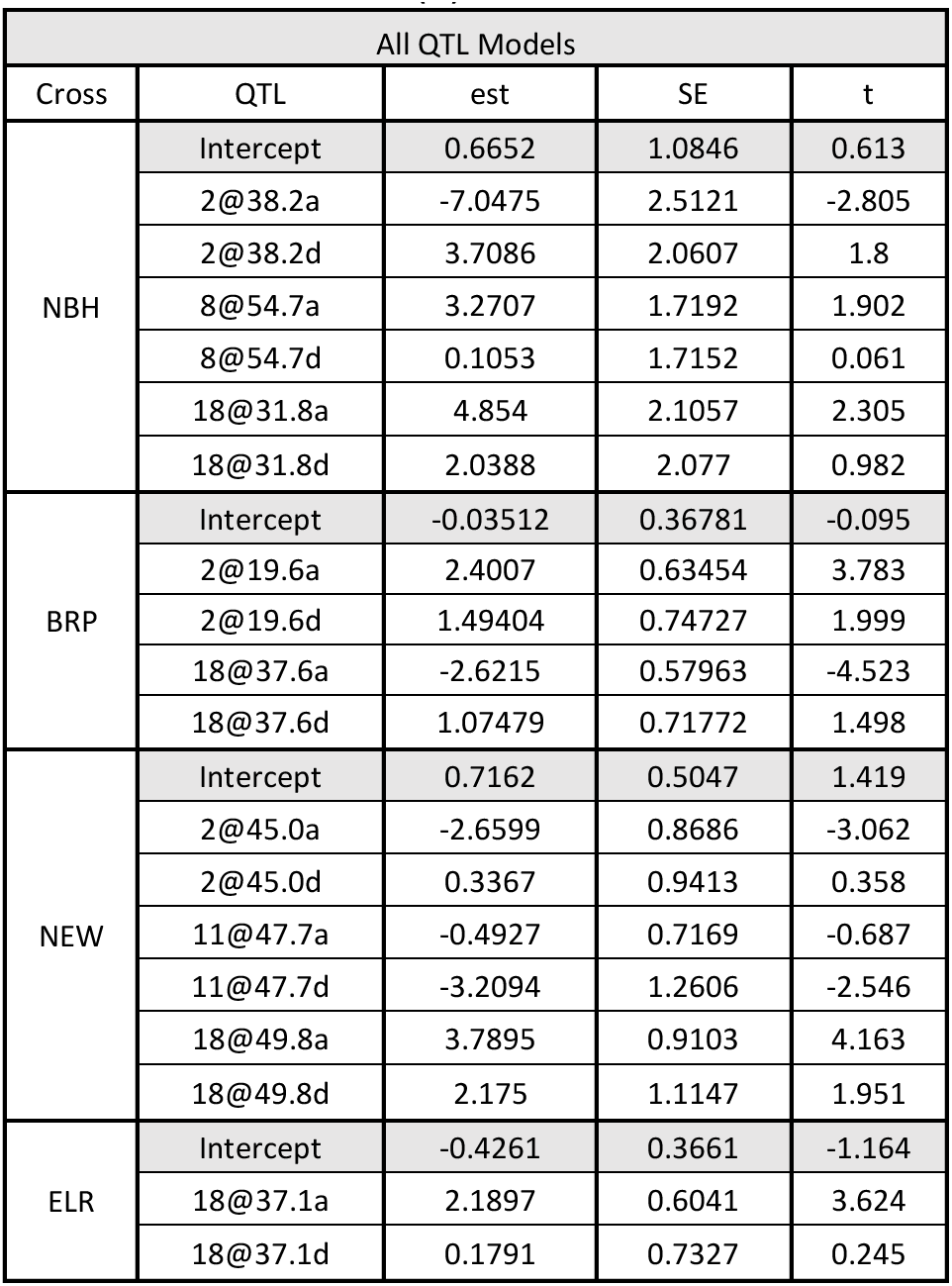
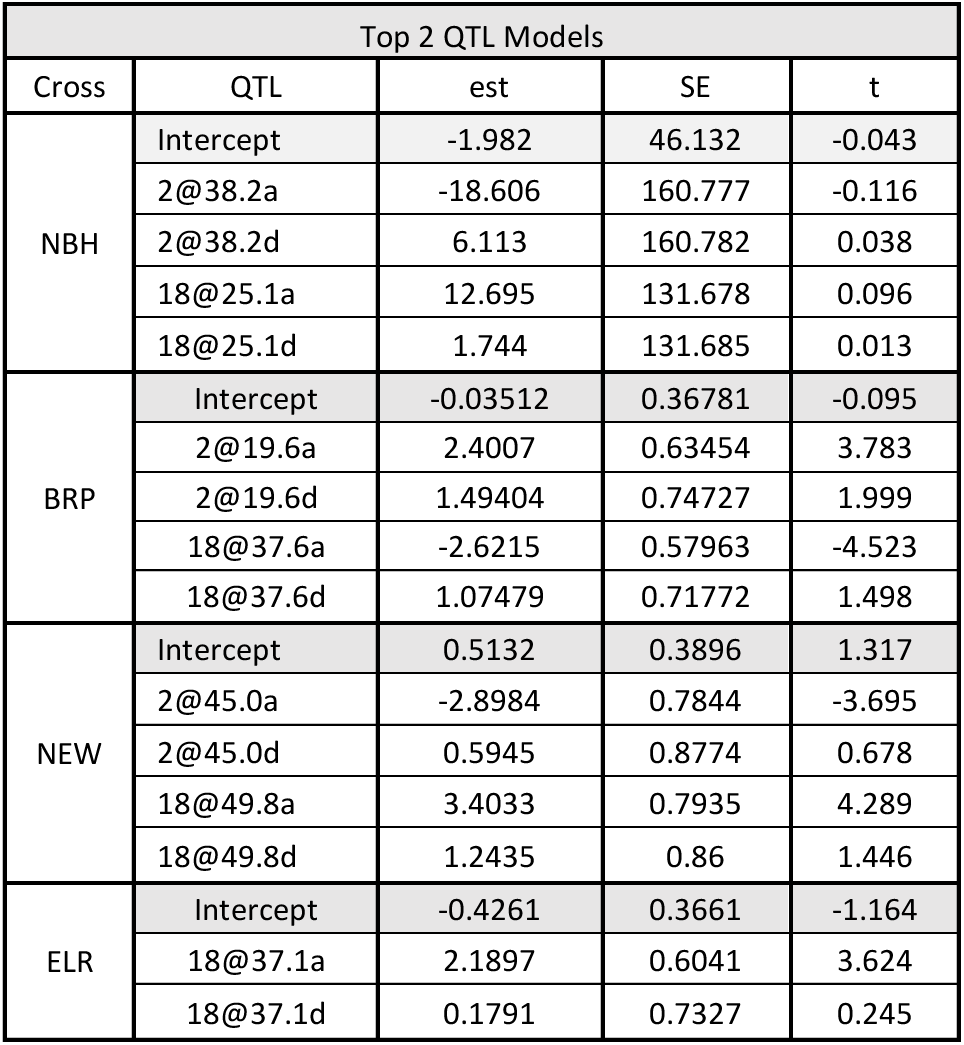
Parameter estimates for QTLs under a binary phenotype using the Haley-Knott regression method in rQTL. Top table of models (All QTL Models) includes estimates for small effect QTL in the full multi-QTL model. Bottom table of models are estimates for each QTL including only the top two QTL in each family included in the full-model. Each QTL in the model is estimated with a dominant effect (d) and an additive effect (a).

| All QTL Models |  |  |  |  |
| --- | --- | --- | --- | --- |
| Cross | QTL | est | SE | t |
| NBH | Intercept | 0.6652 | 1.0846 | 0.613 |
|  | 2@38.2a | -7.0475 | 2.5121 | -2.805 |
|  | 2@38.2d | 3.7086 | 2.0607 | 1.8 |
|  | 8@54.7a | 3.2707 | 1.7192 | 1.902 |
|  | 8@54.7d | 0.1053 | 1.7152 | 0.061 |
|  | 18@31.8a | 4.854 | 2.1057 | 2.305 |
|  | 18@31.8d | 2.0388 | 2.077 | 0.982 |
| BRP | Intercept | -0.03512 | 0.36781 | -0.095 |
|  | 2@19.6a | 2.4007 | 0.63454 | 3.783 |
|  | 2@19.6d | 1.49404 | 0.74727 | 1.999 |
|  | 18@37.6a | -2.6215 | 0.57963 | -4.523 |
|  | 18@37.6d | 1.07479 | 0.71772 | 1.498 |
| NEW | Intercept | 0.7162 | 0.5047 | 1.419 |
|  | 2@45.0a | -2.6599 | 0.8686 | -3.062 |
|  | 2@45.0d | 0.3367 | 0.9413 | 0.358 |
|  | 11@47.7a | -0.4927 | 0.7169 | -0.687 |
|  | 11@47.7d | -3.2094 | 1.2606 | -2.546 |
|  | 18@49.8a | 3.7895 | 0.9103 | 4.163 |
|  | 18@49.8d | 2.175 | 1.1147 | 1.951 |
| ELR | Intercept | -0.4261 | 0.3661 | -1.164 |
|  | 18@37.1a | 2.1897 | 0.6041 | 3.624 |
|  | 18@37.1d | 0.1791 | 0.7327 | 0.245 |

| Top 2 QTL Models |  |  |  |  |
| --- | --- | --- | --- | --- |
| Cross | QTL | est | SE | t |
| NBH | Intercept | -1.982 | 46.132 | -0.043 |
|  | 2@38.2a | -18.606 | 160.777 | -0.116 |
|  | 2@38.2d | 6.113 | 160.782 | 0.038 |
|  | 18@25.1a | 12.695 | 131.678 | 0.096 |
|  | 18@25.1d | 1.744 | 131.685 | 0.013 |
| BRP | Intercept | -0.03512 | 0.36781 | -0.095 |
|  | 2@19.6a | 2.4007 | 0.63454 | 3.783 |
|  | 2@19.6d | 1.49404 | 0.74727 | 1.999 |
|  | 18@37.6a | -2.6215 | 0.57963 | -4.523 |
|  | 18@37.6d | 1.07479 | 0.71772 | 1.498 |
| NEW | Intercept | 0.5132 | 0.3896 | 1.317 |
|  | 2@45.0a | -2.8984 | 0.7844 | -3.695 |
|  | 2@45.0d | 0.5945 | 0.8774 | 0.678 |
|  | 18@49.8a | 3.4033 | 0.7935 | 4.289 |
|  | 18@49.8d | 1.2435 | 0.86 | 1.446 |
| ELR | Intercept | -0.4261 | 0.3661 | -1.164 |
|  | 18@37.1a | 2.1897 | 0.6041 | 3.624 |
|  | 18@37.1d | 0.1791 | 0.7327 | 0.245 |

## Supplemental Files

File S1. binary_model_output.docx

