## Supplementary material for "Independently evolved pollution resistance in four killifish populations is largely explained by few variants of large effect": File S1

NBH Binary (all)

> summary(fit3_hk_bin_perm)

fitqtl summary

Method: Haley-Knott regression

Model: binary phenotype

Number of observations : 91

Full model result

----------------------------------

Model formula: y ~ Q1 + Q2 + Q3

df	LOD	%var Pvalue(Chi2)

Model 6 25.27854 72.17549	0

Drop one QTL at a time ANOVA table:

----------------------------------

df	LOD %var Pvalue(Chi2)

2@38.2	2 17.544 39.79	< 2e-16 ***

8@54.7	2 1.563 2.29	0.0274 *

18@31.8 2 6.817 11.46	1.52e-07 ***

---

Signif. codes: 0 ‘***’ 0.001 ‘**’ 0.01 ‘*’ 0.05 ‘.’ 0.1 ‘ ’ 1

Estimated effects:

-----------------

est	SE	t

Intercept 0.6652 1.0846 0.613

2@38.2a	-7.0475 2.5121 -2.805

2@38.2d	3.7086 2.0607 1.800

8@54.7a	3.2707 1.7192 1.902

8@54.7d	0.1053 1.7152 0.061

18@31.8a	4.8540 2.1057 2.305

18@31.8d	2.0388 2.0770 0.982

BRP Binary

> summary(fit_hk_bin_perm )

fitqtl summary

Method: Haley-Knott regression

Model: binary phenotype

Number of observations : 91

Full model result

----------------------------------

Model formula: y ~ Q1 + Q2

df	LOD	%var Pvalue(Chi2)

Model 4 14.98096 53.14584 3.708145e-14

Drop one QTL at a time ANOVA table:

----------------------------------

df	LOD %var Pvalue(Chi2)

2@19.6	2 7.479 21.55	3.32e-08 ***

18@37.6 2 9.225 27.88	5.96e-10 ***

---

Signif. codes: 0 ‘***’ 0.001 ‘**’ 0.01 ‘*’ 0.05 ‘.’ 0.1 ‘ ’ 1

Estimated effects:

-----------------

est	SE	t

Intercept -0.03512 0.36781 -0.095

2@19.6a	2.40070 0.63454 3.783

2@19.6d	1.49404 0.74727 1.999

18@37.6a -2.62150 0.57963 -4.523

18@37.6d	1.07479 0.71772 1.498

NEW Binary (all)

> summary(fit_hk_bin_perm)

fitqtl summary

Method: Haley-Knott regression

Model: binary phenotype

Number of observations : 85

Full model result

----------------------------------

Model formula: y ~ Q1 + Q2 + Q3

df	LOD	%var Pvalue(Chi2)

Model 6 17.26305 60.75262 4.551914e-15

Drop one QTL at a time ANOVA table:

----------------------------------

df	LOD	%var Pvalue(Chi2)

2@45.0	2 3.672 8.638	0.000213 ***

11@47.7 2 2.332 5.284	0.004661 **

18@49.8 2 12.337 37.330	4.6e-13 ***

---

Signif. codes: 0 ‘***’ 0.001 ‘**’ 0.01 ‘*’ 0.05 ‘.’ 0.1 ‘ ’ 1

Estimated effects:

-----------------

est	SE	t

Intercept 0.7162 0.5047 1.419

2@45.0a	-2.6599 0.8686 -3.062

2@45.0d	0.3367 0.9413 0.358

11@47.7a -0.4927 0.7169 -0.687

11@47.7d -3.2094 1.2606 -2.546

18@49.8a	3.7895 0.9103 4.163

18@49.8d	2.1750 1.1147 1.951

ELR Binary

> summary(fit_hk_bin_perm )

fitqtl summary

Method: Haley-Knott regression

Model: binary phenotype

Number of observations : 59

Full model result

----------------------------------

Model formula: y ~ Q1

df	LOD	%var Pvalue(Chi2)

Model 2 5.362445 34.20056 4.340651e-06

Estimated effects:

-----------------

est	SE	t

Intercept -0.4261 0.3661 -1.164

18@37.1a	2.1897 0.6041 3.624

18@37.1d	0.1791 0.7327 0.245

NBH Binary Large effect QTL only

> summary(fit2_hk_bin_perm)

fitqtl summary

Method: Haley-Knott regression

Model: binary phenotype

Number of observations : 91

Full model result

----------------------------------

Model formula: y ~ Q1 + Q2

df	LOD	%var Pvalue(Chi2)

Model 4 24.15208 70.54324	0

Drop one QTL at a time ANOVA table:

----------------------------------

df	LOD %var Pvalue(Chi2)

2@38.2	2 16.988 40.13	< 2e-16 ***

18@25.1 2 6.252 10.96	5.6e-07 ***

---

Signif. codes: 0 ‘***’ 0.001 ‘**’ 0.01 ‘*’ 0.05 ‘.’ 0.1 ‘ ’ 1

Estimated effects:

-----------------

est	SE	t

Intercept -1.982 46.132 -0.043

2@38.2a	-18.606 160.777 -0.116

2@38.2d	6.113 160.782 0.038

18@25.1a	12.695 131.678 0.096

18@25.1d	1.744 131.685 0.013

NEW Binary Largest effect QTL only model

> summary(fit2_hk_bin_perm)

fitqtl summary

Method: Haley-Knott regression

Model: binary phenotype

Number of observations : 85

Full model result

----------------------------------

Model formula: y ~ Q1 + Q2

df	LOD	%var Pvalue(Chi2)

Model 4 14.93155 55.46828 4.141132e-14

Drop one QTL at a time ANOVA table:

----------------------------------

df	LOD %var Pvalue(Chi2)

2@45.0	2 6.286 18.07	5.18e-07 ***

18@49.8 2 11.888 40.27	1.29e-12 ***

---

Signif. codes: 0 ‘***’ 0.001 ‘**’ 0.01 ‘*’ 0.05 ‘.’ 0.1 ‘ ’ 1

Estimated effects:

-----------------

est	SE	t

Intercept 0.5132 0.3896 1.317

2@45.0a	-2.8984 0.7844 -3.695

2@45.0d	0.5945 0.8774 0.678

18@49.8a	3.4033 0.7935 4.289

18@49.8d	1.2435 0.8600 1.446
